# Methylglyoxal controls tomato fruit ripening by regulating ethylene biosynthesis

**DOI:** 10.1101/2022.11.03.515062

**Authors:** Priya Gambhir, Utkarsh Raghuvanshi, Adwaita Prasad Parida, Stuti Kujur, Shweta Sharma, Sudhir K. Sopory, Rahul Kumar, Arun Kumar Sharma

## Abstract

Methylglyoxal (MG), a toxic compound produced as a byproduct in several cellular processes such as respiration and photosynthesis, is well investigated for its deleterious effects, mainly through glycation of proteins during plant stress responses. However, very little is known about its impact on fruit ripening. In the present study, we report that MG levels are maintained at high level in green tomato fruits, which declines during fruit ripening inspite of a respiratory burst during this transition. We demonstrate that this decline is mainly mediated by glutathione-dependent MG detoxification pathway and primarily catalyzed by glyoxalase enzyme encoded by *SlGLY14* gene. *SlGLYI4* is a direct target of MADS-RIN and is induced during fruit ripening. Silencing of this gene leads to drastic MG overaccumulation at ripening-stages in the transgenic fruits and interferes with the ripening process. Further investigations show that MG plausibly glycates and inhibits key enzymes such as methionine synthase (MS) and S-adenosyl methionine synthase (SAMS) of ethylene biosynthesis pathway, thereby indirectly affecting fruit pigmentation and cell was metabolism. MG overaccumulation in several non-ripening or inhibited- ripening tomato mutant fruits suggests the tightly regulated MG detoxification process is crucial for normal ripening program. Overall, we underpin a *SlGLYI4-*mediated novel regulatory mechanism of MG detoxification controlling fruit ripening in tomato.

## Introduction

Fruit ripening is a multifactorial process often paralleled to a functionally modified protracted form of senescence encompassing ethylene burst, membrane remodeling, and alteration in levels of pigments with a concomitant increase in total, reducing, and non-reducing sugars (Jimenez *et al*., 2002; Anwar *et al*., 2018; Figueroa *et al*., 2021). This inevitably creates a state of oxidative stress when the fruits start to mature (Sharma *et al*., 2009). These alterations in fruit’s oxidative metabolism serve two separate but very significant roles: regulating fruit’s growth and architecture or forming an essential trait for optimizing biochemical and genetic reprogramming necessary for the progression of climacteric and non-climacteric ripening.

To date, oxidative metabolism during tomato fruit ripening has majorly focused on reactive oxygen species (ROS) and reactive nitrogen species (RNS) equilibrium (Jimenez *et al*., 2002; Mondal *et al*., 2004; Kumar *et al*., 2016; Corpas *et al*., 2018). The maiden reports on an intensified defense mechanism against ROS synthesis in photosynthetically active immature green tomato fruits dates to the early 2000s, with studies demonstrating enrichment of the CAT, SOD, MDAR, DHAR, and GR enzyme activities during fruit growth and maturation (Jimenez *et al*., 2002). The conversion of chloroplast into chromoplast via the dismantling of thylakoids strongly influences fruit ripening.

During this transition, photosynthesis-mediated ROS production is taken over by mitochondrial respiration-mediated ROS synthesis in fruits undergoing ripening (Maxwell *et al*., 1999). It resonates perfectly well with the reported transitory rise in H_2_O_2_ levels amalgamated with a high lipid/protein oxidation content in tomato fruits (Rabinowitch *et al*., 1982; Jiménez *et al*., 2002; Kumar *et al*., 2016). Numerous studies have hypothesized that a rise in oxidants production is directly proportional to the initiation and progression of tomato fruit ripening, as exemplified by the substantially low levels of ROS in the fruits and failure of attainment of ripening in mutants such as *rin* and *nor* (Mehta *et al*., 2002; Mondal *et al*., 2004; Zhang *et al*., 2013; Kumar *et al*., 2016). The fugacious rise in ROS levels witnessed at the onset of fruit ripening is counterbalanced by the activation of ascorbate and glutathione biosynthesis pathways (Jimenez *et al*., 2002; Mondal *et al*., 2004; Murshed *et al*., 2014). The ratio of the reduced glutathione to the total glutathione and the enzymatic activity of glutathione reductase (GR) responsible for the glutathione redox state has been shown to gradually amplify throughout ripening (Jiménez *et al*., 2002; Corpas *et al*., 2018). Hence, the reinforcement of the antioxidant defense mechanism in fully ripe tomato fruits culminates in an enhanced accumulation of antioxidants such as carotenoids, flavonoids, ascorbate, and tocopherols, thereby managing the oxidative stress perceived at the final stages of ripening. This extreme upregulation in ascorbate-dependent ROS biosynthesis has been suggested to evoke a series of response imperative for destabilizing the plant cell wall in a non-enzymatic way to tomato fruit softening (Jimenez *et al*., 2002; Dumville and Fry, 2003). The production and accumulation of carotenoids and phenylpropanoids in tomato fruits have also been shown to be directly or indirectly regulated by the cellular redox state (Fanciullino *et al*., 2014). Interestingly, the exogenous application of some compounds such as H_2_O_2_, nitric oxide (NO), and melatonin imparted various beneficial effects to fruits, as evidenced by delayed senescence and enhanced nutritional aspect. Their application also alleviates fungal decay of the ripened fruits via employing the enzymatic and non-enzymatic antioxidant systems (Manjunatha *et al*., 2012; Bodanapu *et al*., 2016; Zhou *et al*., 2016; Liu *et al*., 2019). Engineering tomato fruits for enhanced anthocyanins and antioxidant levels also prolong fruit longevity via delayed senescence and compromised ROS production (Zhang *et al*., 2013). Likewise, transgenic tomatoes with high spermine and spermidine content, two polyamines tagged with pivotal antioxidant properties, resulted in improved lycopene accumulation and fruit shelf-life (Mehta *et al*., 2002).

The role of NO and RNS as potential regulators of fruit ripening has also emerged (Leshem *et al*., 1998; Leshem and Pinchasov, 2000). Like ROS, RNS equilibrium is significantly altered as the fruits start to ripen (Jiménez *et al*., 2002; Kumar *et al*., 2016; Corpas *et al*., 2018). NO has been in the limelight as a promising molecule to prolong the postharvest shelf life without impacting the color, aroma, texture, and nutritional value of fruits (Manjunatha *et al*., 2012; Corpas *et al*., 2018; Zuccarelli *et al*., 2021). External NO application was found to promote enhanced accumulation of phenolics in ripe peach and citrus (Zhou *et al*., 2016; Li *et al*., 2017). Accumulating evidence points towards a defective ethylene biosynthetic pathway as one of the primary outcomes of enhanced RNS accumulation during climacteric fruit ripening (Rudell and Mattheis, 2006; Zhu and Zhou, 2006, 2007; Eum *et al*., 2008; Cheng *et al*., 2009; Palma *et al*., 2019). This off-balance modulates the entire arena of the fruit’s oxidative state with specific ramifications on NO-mediated post-translational modifications of target proteins (Chaki *et al*., 2015; Corpas *et al*., 2018; Palma *et al*., 2019).Thus, NO is often reckoned as a two-way sword with its positive influence on the transcript levels of genes involved in antioxidant (enzymatic and non-enzymatic) defenses, whereas targeting the same antioxidant enzymes for nitration and S-nitrosation in the fruits (Groß *et al*., 2013; Boscari *et al*., 2013; Chaki *et al*., 2015; Huang *et al*., 2018; Corpas *et al*., 2018; Rodríguez-Ruiz *et al*., 2019; González-Gordo *et al*., 2019; Palma *et al*., 2019; Kolbert *et al*., 2019).

Despite substantial progress toward understanding the ripening-associated changes in fruit ROS and RNS metabolism, our knowledge of all the oxidative state contributors at the ripening onset is far from complete. This juxtaposes with the immense agronomical value that is bestowed upon the tomato as a model species for cognizing fleshy fruit development and climacteric mode of ripening. Like ROS and RNS, one such molecule, Methylglyoxal (MG), is implicated as a marker of oxidative stress but, interestingly, has never been studied in the context of fruit ripening (Hoque *et al*., 2016a). MG is a by- product of many physiological processes targeting plant growth and differentiation (Thornalley, 1990; Phillips and Thornalley, 1993; Yadav *et al*., 2005; Takagi *et al*., 2014; Kaur *et al*., 2015). It was initially discovered in plants in 1990 as a compound with no known metabolic function. However, its overaccumulation was found to terminate in deleterious events, ultimately leading to cell death (Thornalley, 1990; Phillips and Thornalley, 1993). Since then, MG has been constantly scrutinized for its function as an intensifier of ROS synthesis and a powerful cell growth inhibitor in numerous plant species (Guo *et al*., 2016; Hoque *et al*., 2016*a*; Li, 2016; Rabbani, 2022). MG is a highly potent glycating agent and readily interacts with nucleic acids, proteins, and lipids to create advanced glycation end products (Thornalley, 1990; Asada, 1996; Hoque *et al*., 2016*b*; Chaplin *et al*., 2019). Besides being cytotoxic, MG has emerged as a critical regulator of stomatal conductance, encompassing cytosolic Ca^+^ oscillations and potassium channel activation (Hoque *et al*., 2012*a*,*c*, 2016*b*). The elevated level of MG is instrumental when the plants are challenged with multiple abiotic stress conditions, including salt, osmotic and oxidative stress (Hoque *et al*., 2016*b*; Gupta *et al*., 2018). Two glyoxalase detoxification pathways are known to maintain and reinforce cellular MG equilibrium. The first pathway is a glutathione-dependent bi-enzymatic glyoxalase system. This pathway has received renewed interest for catalyzing the conversion of methylglyoxal to D-lactate via the production of an intermediate product, S-(D-lactoyl) glutathione, and sustaining glutathione (GSH) repertoire upon encountering stressful circumstances. The second pathway for MG removal from the cell is shorter, more direct, and independent of GSH. It is catalyzed by the recently identified Glyoxalase III class of enzymes (Ghosh *et al*., 2016).

In this study, we have investigated the impact of MG accumulation on tomato fruit ripening. Our findings indicated a key role of MG detoxification concerning the progression of tomato fruit ripening. We also deciphered the putative mechanism of how MG possibly tempers with fruit ripening. For this purpose, we generated artificial microRNA lines of a ripening-associated glyoxalase gene, *SlGLYI4*, actively involved in fruit growth and maturation under two different sets of promoters, the *CaMV35S* and *RIP (Ripening-induced protein 1)* (Agarwal *et al*., 2017). A comprehensive analysis of morphological, biochemical, and physiological parameters of *35S:amiRGLYI4* and *RIP:amiRGLYI4* silencing lines revealed that elevated MG levels inhibit fruit ripening by inhibiting ethylene biosynthesis, plausibly glycating the enzymes involved in methionine and 1-aminocyclopropane-1-carboxylic acid (ACC) and inhibiting their accumulation. In summary, we demonstrate the significance of MG levels and its detoxification process on the overall development of fruit and the ripening process.

## Materials and Methods

### Plant materials and growth conditions

Tomato (*Solanum lycopersicum* L.cv Pusa Ruby) WT, *35S:RIN-RNAi* transgenic lines in Pusa Ruby background generated in our lab (RIAI05 and RIAI06) (http://hdl.handle.net/10603/389794), Ailsa Craig WT and *ripening-inhibitor* (*rin*, accession no. LA1795, in an unknown background)*, non-ripening* (*nor*, Ailsa Craig background) and Moneymaker WT and *high-pigment 2* (*hp2*, Moneymaker background) fruit ripening mutants were grown under standard greenhouse conditions: 14-h-day/10-h-night cycle, 25°C/20°C day/night temperature, 60% relative air humidity, and 250 µmol m^−2^ s^−1^ intense luminosity. Fruit pericarp tissue samples were collected from different fruit development and ripening stages, as described earlier (Kumar *et al*., 2012). For the measurement of time to ripen, flowers were tagged at anthesis. The number of days counted from anthesis to the appearance of the first symptoms of ripening was designated as the number of days taken to reach the breaker (Br) stage. All fruit stages were harvested in five biological replicates. Fruits from each genotype were harvested at different stages and, fruits were snap-frozen in liquid nitrogen to avoid injury.

### Treatment to fruits

#### MG treatment

For MG treatment, tomato fruits either on vine (MT background) or harvested (Pusa Ruby background) at the mature green stage were injected with 10 or 25 mM aqueous solution of MG (Sigma) using a 1 ml syringe with a 0.5 mm needle and inserted 3 to 4 mm into the fruit tissue through the stylar apex. The treatment was repeated two days after the first injection. After the treatment, the harvested fruits were incubated in a culture room at 26°C, under a 16 h light/8 h dark cycle with a light intensity of 100 μmol m^-2^s^-1^. Both on-vine and the harvested fruits were monitored until the attainment of ripening. Distilled water-infiltrated fruits acted as the control. Pericarp tissue of the treated fruits was collected and frozen in liquid nitrogen for further biochemical analysis.

#### Ethylene, 1-MCP, and ACC treatment

Exogenous application of ethylene and 1-MCP was done as previously described (Gambhir *et al*., 2022). Briefly, for ethylene and ACC treatment, wild type tomato fruits (Pusa Ruby) were harvested at the mature green stage and injected with a buffer solution containing 10 mM MES, pH 5.6, sorbitol (3% w/v), and 100 μM of Ethrel (2-Chloroethylphosphonic Acid, 40% Solution, Sisco Research Laboratories Pvt. Ltd. India) or 100 μM of ACC respectively. For ethylene inhibitor treatment, tomato fruits harvested at mature green stage were infiltrated with the abovementioned buffer containing 100 µM 1-MCP.

The infiltration solution was gently injected into the fruit until the solution ran off the stylar apex. Only completely infiltrated fruits were used in the experiments. Control fruits were treated with the corresponding buffers only. The treated fruits were incubated in a culture room at 26°C, under a 16 h light/8 h dark cycle with a light intensity of 100 μmol m^-2^s^-1^. After 24 h, the fruit pericarp was collected and frozen at -80°C until further use.

#### Methionine treatment

Pusa Ruby fruits harvested at the mature green stage were infiltrated with 10 mM methionine (Sisco Research Laboratories Pvt. Ltd. India). After treatment, the fruits were kept at 26°C, under a 16 h light/8 h dark cycle with a light intensity of 100 μmol m^-2^s^-1^ overnight, then adequately sealed in glass vials provided with a rubber septum. 1-ml of the headspace gas was further analyzed for ethylene measurement assay.

### RNA isolation and quantitative real-time polymerase chain reaction (qRT- PCR)

Total RNA from tomato leaves and fruitpericarp tissues was isolated using RNeasy Plant Mini Kit (Qiagen, Germany) following the manufacturer’s instructions. The provision for on-column DNase-treatment was provided with the kit. 1-µg of total RNA for each sample was subjected to cDNA synthesis with high-capacity cDNA reverse transcription kit (Applied Biosystems, USA). Gene-specific primers for qRT-PCR were designed using PRIMER EXPRESS version 2.0 (PE Applied Biosystems, USA) with default parameters. Further, 2X Brilliant III SYBR® Green QPCR master mix (Agilent Technologies, USA) was used for a qRT-PCR reaction carried out in a Stratagene Mx3005P qPCR machine (Agilent Technologies, USA). Five independent RNA isolations and three technical replicates were used for mRNA quantification. The expression values of genes were normalized *Actin* gene expression values. Relative expression values were calculated using the 2^-ΔΔCT^ method (Livak and Schmittgen, 2001).

### Construction of artificial microRNA-based constitutive and ripening specific *SlGLYI4* silencing vectors and tomato transformation

The coding sequence of *SlGLYI4* was used as an input query for the wmd2-web microRNA designer platform (http://wmd2.weigelworld.org) to retrieve amiRNA sequence. A few amiRNAs starting with uridine residue and targeting different regions of *SlGLYI4* were manually selected and screened for the absence of any off-targets in tomato genome. Finally, amiRNA 5′ UUCGUUAAUCCUGGCAUGCUU 3′ targeting the 1323-1343 nt region of *SlGLYI4* mRNA was selected for cloning. *Arabidopsis thaliana* pre- miRNA319a backbone was chosen for expression of *SlGLYI4*-amiR in transgenic tomato. Recombinant *SlGLYI4*-preamiRNA1 was PCR amplified by extension of overlapping primers (Supporting table S1), ligated onto the pUC19 vector, and sequence confirmed. Then *SlGLYI4*-preamiRNA1 was excised from the carrier plasmid and ligated between the *CaMV35S* promoter and NOS terminator of plant binary vector, pBI121 to get the amiRNA vector cassette, *35S:amiRGLYI4.* Similarly*, SlGLYI4*-preamiRNA1 cassete was cloned under the RIP (Ripening-specific) promoter in the modified pBI121 (Agarwal *et al*., 2017) to get *RIP1:amRGLYI4 vector* constructs. The amiR plasmids were mobilized from *E. coli* into *Agrobacterium* tumefaciens strain AGL1. After confirmation of this mobilization, the transformed *Agrobacterium* cultures were used to generate stable transgenic lines of *35S:amiRGLYI4* and *RIP1:amiRGLYI4* in Pusa Ruby, as described previously (Sharma *et al*., 2009). Primers used in constructing *35S:amiRGLYI4* and *RIP1:amiRGLYI4* constructs are listed in Supporting Table S1. Both sets of amiR-based *SlGLYI4*-silenced lines were grown and analyzed for their morphological, biochemical, molecular, and physiological characterization in the T_2_ generation. The segregation analysis of kanamycin resistance in T_2_ progeny of *35S:amiRGLYI4* and *RIP1:amiRGLYI4* transgenic tomato plants was done to obtain homozygous lines (Supporting Table S2). Final experiments were carried out using homozygous lines from T_2_ or later generations.

### Virus-induced gene silencing

Tobacco rattle virus (TRV)-based *pTRV1* and *pTRV2* vectors were used for Virus-induced gene silencing (VIGS) experiments (Liu *et al*., 2002). A 300-bp fragment of the coding region corresponding to *SlGLYI4, SlGLYI7,* and *SlGLYII2* was retrieved by us using the VIGS tool of the SOL Genomics Network Database (https://vigs.solgenomics.net/). Each 300-bp fragment was then PCR amplified from Pusa Ruby cDNA and inserted in the *pTRV2* vector using *Xba1* and *EcoR1* restriction sites for *SlGLYI4* and *SlGLYI7* and *Xho1* and *Sac1* for *SlGLYII2*. The primers used for VIGS assay cloning are listed in the Supporting Table S1. The empty *pTRVs* vector was used as a control in the VIGS assays. The *pTRV::GLYI4*, *pTRV::GLYI7,* and *pTRV::GLYII2* plasmids verified by sequencing were then mobilized into *Agrobacterium tumefaciens* strain GV3101. VIGS was carried out on mature green fruits (36DPA), as described previously (Fu *et al*., 2005).

### Measurement of enzyme activities

For enzyme assays, protein extract was prepared by homogenizing pericarp tissue from WT controls, ripening mutants (*rin*, *nor* and *hp2*), and *SlGLYI4*- silenced transgenic plants at different stages of ripening in liquid nitrogen. Powdered pericarp homogenates were resuspended in 2 vol (w/vol) of extraction buffer (0.1 M sodium phosphate buffer, pH 7.0/ 50% glycerol/16 mM MgSO_4_/0.2 mM PMSF/ 0.2% polyvinyl polypyrrolidone). The concentration of the protein present in the crude extract was measured spectrophotometrically using the Bradford method (Ku *et al*., 2013).

#### GLY1 and GLYII activity

The enzyme activity for GLYI and GLYII was determined as described earlier (Zeng *et al*., 2016). Briefly, the crushed tissue extract was centrifuged twice at 13,000 rpm at 4°C for 30 min to obtain the protein portion from the supernatant. GLYI and GLYII activity was assayed using an extinction coefficient of 3.37 mM^−1^ cm^−1^. Three different enzyme extractions were done per sample for three independent transgenic lines. The specific activity of both enzymes is expressed in units mg^-1^ of protein.

#### PL activity

The enzymatic activity of pectate lyase was assayed as previously described (Collmer *et al*., 1988). Briefly, fresh pericarp tissue (Br+8) was homogenized in the presence of ice-cold 80% (v/v) acetone for the preparation of acetone insoluble solids (AIS). The homogenized sample was washed with 100% acetone to eliminate all the pigments. Afterward, the powder was left to dry overnight at room temperature. Then 2.5 mg of the purified AIS was stirred in 0.95 ml of 8.5 M Tris-HCL at 20^◦^C for approximately 30 min. The samples were then subjected to centrifugation at 8,000 g for 30 min, and the absorbance of clear supernatant was quantified at 232 nm to determine the extent of reaction products with double bonds released as a result of PL activity. Control assays were conducted where AIS was inactivated by boiling in 80% (v/v) ethanol.

### Determination of MG, D-Lactate, GSH/GSSG ratio, MDA, and total antioxidant content

Pericarp tissue was collected from the WT controls, ripening mutants (*rin*, *nor* and *hp2*) and transgenic fruits at different stages of ripening. MG content was measured as described previously (Yadav *et al*., 2005). GSH/GSSG ratio was calculated using a glutathione GSH/GSSG colorimetric assay kit (Sigma-Aldrich, USA). For D-lactate measurement, 50 μl of the neutralized extract was added to the reaction mixture containing 100 μl of potassium phosphate buffer, 40 μl of 1 mM DCIP, 10 μl of 60 mM PMS and 1 μl of enzyme extract (0.025 U/μl). The change in absorbance was recorded at 600 nm for 5 min taking a reading at an interval of 30s. D-lactate content was calculated using a standard curve. For the standard curve, different concentrations of D-lactate (0–400 nmole) were used and assayed similarly. Malondialdehyde (MDA) content, a measure of lipid peroxidation, was measured by thiobarbituric acid extraction protocol, as described previously (Boaretto *et al*., 2014). Total antioxidant content was determined from pericarp tissue of WT and transgenic lines by ferric reducing antioxidant power (FRAP) assay following the standard protocol, as previously described (Wong *et al*., 2006).

### Fruit firmness measurement

The assessment of fruit firmness was carried out using TA.XT Plus Texture Analyser (Stable Micro Systems, England) as previously described (Gambhir *et al*., 2022). Briefly, fruits of wild type and SlGLYI4-silenced genotypes during different stages of ripening stage were subjected to a puncture test with a 2 mm needle probe, and the force and distance measurements were recorded. Tenfruits from each transgenic and tissue culture generated WT genotype were used in the study. Fruit firmness calculated is equivalent to the amount of force applied to penetrate the surface of the fruit.

### Ethylene measurement

Ethylene measurement was done as previously described (Gambhir *et al*., 2022). Briefly, fruits at different ripening stages were harvested and stored in open 250-ml jars for 2 h to minimize the effect of wound-induced ethylene caused by the harvesting of the fruits. Jars were then sealed and incubated at room temperature for 4 h, and 1 ml of headspace gas was injected into Shimadzu QP-2010 Plus with Thermal Desorption System TD 20 (Shimadzu, Japan). Samples were compared with reagent-grade ethylene standards of known concentration and normalized for fruit weight. Ethylene in the headspace gas was measured thrice for each sample, with at least five biological replicates for each ripening stage.

### Estimation of fruit pigments

Lycopene and β-carotene extractions for HPLC experiments were performed as described previously (Gambhir *et al*., 2022). Briefly, 150 mg of ground lyophilized tomato fruit powder were extracted with chloroform and methanol (2:1 v/v); Then, 1 volume of 50 mM Tris buffer (pH 7.5, containing 1M NaCl) was added followed by the incubation of samples on ice for 20 min. After centrifugation (15,000 *g* for 10 min at 4°C), the organic phase was collected, and the aqueous phase was re-extracted with the same amount of chloroform. The combined organic phases were then dried by centrifugal evaporation and resuspended in 100 µl of ethyl acetate. A final volume of 20 µl was injected into a C-18 column in HPLC (Shimadzu, Japan) analysis. For each genotype, at least five independent extractions were performed.

### Pectin measurement

For protopectin and soluble pectin estimation, 500 mg of tomato pericarp tissue (devoid of any placental portions) obtained from WT and transgenic lines at the Br+8 stage was homogenized with 25 ml of 90% ethanol and boiled for about 30 min. After cooling at room temperature, the samples were subjected to centrifugation at 8,000 *g* for 15 min. Subsequently, the supernatant was removed, and the pellet was again resuspended in 25 ml of 90% ethanol. This step was repeated 3-4 times, and the final pellet was resuspended in 20 ml of distilled water. The samples were then incubated for 30 min at 50°C. This step was followed by centrifugation at 8,000 *g* for 15 min. The obtained pellet was dissolved in 25 ml of 0.5mol/l of H_2_SO_4_ and boiled for 1h. The samples were again subjected to centrifugation at 8,000 *g* for 15 min, and the supernatant containing the protopectin was retrieved. The pectin content was then measured by mixing 1ml of the collected pectin with 6 ml of concentrated HCl, boiling for 20 min. After cooling at room temperature, 0.2 ml 1.5g l^−1^ carbazole was added to the sample and incubated in the dark at RT for 1 h. The absorbance at 530 nm was measured against reagent blanks, and pectin content was calculated based on a galacturonic acid standard curve.

### Yeast-one-hybrid assay

For yeast-one-hybrid (Y1H) experiments, 1.5 kb long promoter sequence, upstream of the start codon, of *SlGLYI4* was amplified using Pusa Ruby genomic DNA. The amplified products were cloned into the yeast expression vector *pHIS2*.1 and co-transformed with the *pGAD-T7-RIN* into yeast strain *Y187*. The binding of RIN to *LeACS2 (1-aminocyclopropane-1-carboxylate synthase 2)* promoters was taken as positive control respectively, for this experiment. The DNA-protein interaction was validated by transformant growth assays on SD/-Leu/-Trp/-His plates supplemented with 50 mM of 3-AT. Primers used in this section are listed in Supporting Table S1.

### Transactivation of *SlGLYI4* promoter in *N. benthamiana*

To verify the DNA-protein interaction in planta, 1.5 kb upstream region of *SlGLYI4* was used to drive the expression of GFP, designated as a reporter vector. For the effector vector, the full open reading frame (ORF) of *RIN* was cloned in the binary vector *pBI121* driven by the *CaMV35S* promoter. Both reporter and effector constructs were co-transformed into 4-week-old *N. benthamiana* leaves, as described previously (Gambhir *et al*., 2022). GFP fluorescence measurement assay using was done after 48 h of infiltrations. The excitation for GFP was carried out by argon laser, and the fluorescence was detected using a bandpass filter (530/30nm) in the FL1 channel. The NightSHADE LB 985 (Berthold Technologies USA) *in vivo* plant imaging system was employed to detect fluorescent signals with 5s exposure time. Data were analyzed by the IndiGO™ software. The average fluorescence signal for each sample was collected (cps, count per second). It was normalized using the *N. benthamiana* leaves transformed with a reporter vector combined with the vector used as an effector but lacking the *SlGLYI4* or *RIN* coding sequences. Three independent replicates were used for each analysis. Primers used in this experiment are listed in the Supporting table: Table S1.

### Electrophoretic mobility shift assay

The full-length *RIN* coding sequence was used for in-frame cloning into pET-28a (to fuse in-frame with 6X Histidine tag) for heterologous protein expression. The fusion protein construct was then expressed in the BL21 strain of *E. coli*. Recombinant RIN-6XHIS protein was purified by Ni^2+^ gravity flow chromatography according to the manufacturer’s protocol (Ni-NTA Agarose, Qiagen). For the EMSA assay, the 50-bp probe covering the CArG element (CAAAATTAAG) derived from *SlGLYI4* promoter was labelled with digoxigenin as per the manufacturer’s protocol using DIG Oligonucleotide 3′- End Labelling Kit (Roche Diagnostics). The same unlabelled DNA fragment was used as a competitor. The binding reactions were performed at room temperature in binding buffer (10 mM Tris (pH7.5), 50 mM KCl, 1 mM DTT, 2.5% glycerol, 5 mM MgCl_2_, 0.5 mM EDTA, 50 ng/ml poly (dI-dC)) containing 1µg purified RIN-6XHIS fusion protein and 5 nanogram probes. The reaction products were analyzed on 6% (w/v) native polyacrylamide gel electrophoresis. The products were then transferred from the gel to the Hybond N+ Nylon membrane (Amersham Biosciences) and detected using DIG Nucleic Acid Detection Kit (Roche Diagnostics), according to the manufacturer’s protocol.

### Protein extraction and Western blot Analysis

The fruits of Pusa Ruby (WT), *35S:RIN-RNAi* transgenic lines (RIAI05) (http://hdl.handle.net/10603/389794), Ailsa Crag WT and *rin* (accession no. LA1795, in an unknown background)*, nor* (Ailsa Craig background) and *hp2* (Moneymaker background) and *SlGLYI-4*-silenced fruits were collected at different stages of ripening. Total protein extracts were prepared by grinding the fruit pericarp tissue in extraction buffer containing 50 mM Tris-HCl, 50mM NaCl, 10mM MgCl_2_, 5mM EDTA, 5mM DTT, and 10% glycerol (v/v) supplemented with plant protease inhibitor cocktail tablets (Roche, Switzerland). The Bio-Rad protein reagent assay and Coomassie Brilliant Blue R250 staining techniques were employed to determine the protein concentration. Ten micrograms of total protein were separated on 10% SDS-PAGE gel and were followed by protein blot analysis using mouse anti-methylglyoxal monoclonal antibodies (Cell Biolabs, Catalog number- STA-011).

### Determination of free amino acids by UPLC

For analysis of amino acids, 200 mg of fruit pericarp tissue was used. The amino acid content was measured using the protocol described previously (Szkudzińska *et al*., 2017). Briefly, the finely ground test samples were passed through 0.25mm sieve samples and subjected to performic acid oxidation for 16 h in the ice bath. Following oxidation, the sample was hydrolyzed using 6M HCl-phenol solution for 24 hr at 110-120°C. The hydrolysates were then filtered into a 100 ml vacuum flask, and 5ml of norleucine standard solution was added to the sample. 5ml of the samples were evaporated to dryness, and then the dry residue was dissolved in 10 ml of HPLC grade water. The pre-column derivatization was then performed using borate buffer (reagent 1), AQC (reagent 2A), acetonitrile (reagent 2B), and acetonitrile of gradient purity for HPLC. 1-µl of the derivatized samples were then injected into The ACQUITY UPLC system (Waters, Milford, MA, USA) consisting of a thermostat, autosampler, high-pressure binary pump, and photodiode array detector PDA (an optical detector in the range ultraviolet-visible light that operates between 190 nm and 700 nm). Chromatographic separation was obtained with the AccQ-Tag Ultra C-18 column (2.1 mm×100 mm; 1.7 µm).

## Results

### Lower MG level promotes tomato fruit ripening

We first conducted experiments consisting of exogenous application of MG to WT mature green fruits to investigate the role of MG during fruit ripening. Both on-vine or harvested MG-infiltrated fruits exhibited a deep orange to yellow color and contrasted with the red phenotype observed in water-infiltrated control fruits at the Br+8 stage (Fig 1A, Supporting fig, S1C). We next determined the MG concentration in both sets of fruits at the Br+8 stage and noticed a significant enhancement of approximately 55-60% in its levels in MG- infiltrated fruits (10 mM) compared to the water-treated control counterparts (Supporting fig, S1A). Furthermore, given the capacity of MG to interact with proteins and render them non-functional, our next aim was to check the extent of MG-modified proteins in infiltrated and control fruits using an anti-MG antibody. Consistent with the concentrations of MG, the fruits injected with MG unveiled an increase in proteins modified by MG compared to the control fruits (Supporting fig, S1B). Additionaly, the MG-treated fruits exhibited less ethylene production at Br+3 stage as compared to control fruits (Supporting figure, S1D). These results indicate that the failure of the ripening phenotype in MG-treated fruits can be attributed to a possible surge in its levels. Several non-ripening mutants fail to develop red color and *rin* mutant fruits exhibit yellowish color at the ripening-equivalent stage, a phenotype reminiscent of MG-treated fruit. We next profiled fruits of different WT backgrounds and ripening defective mutants i.e., *rin*, *nor* and *hp2* at the mature green, Br, and red ripe (Br+8) stage for MG levels (Fig 1B). This analysis revealed *rin* and *nor* mutant fruits accumulate copious amounts of MG at Br and Br+8 stages, whereas only a slight increase in its amount was observed in *hp2* mutant in comparison to their respective WT fruits (Fig 1B). Correspondingly, higher protein modifications by MG were witnessed in *rin* and *nor* fruits than their corresponding WTs. Interestingly, Moneymaker background fruits (WT and *hp2* mutant) exhibited the least effect of MG on protein modification (Fig 1C). These observations imply that fluctuations in MG equilibrium at ripening stages would adversely impact tomato fruit ripening.

**Figure 1.**
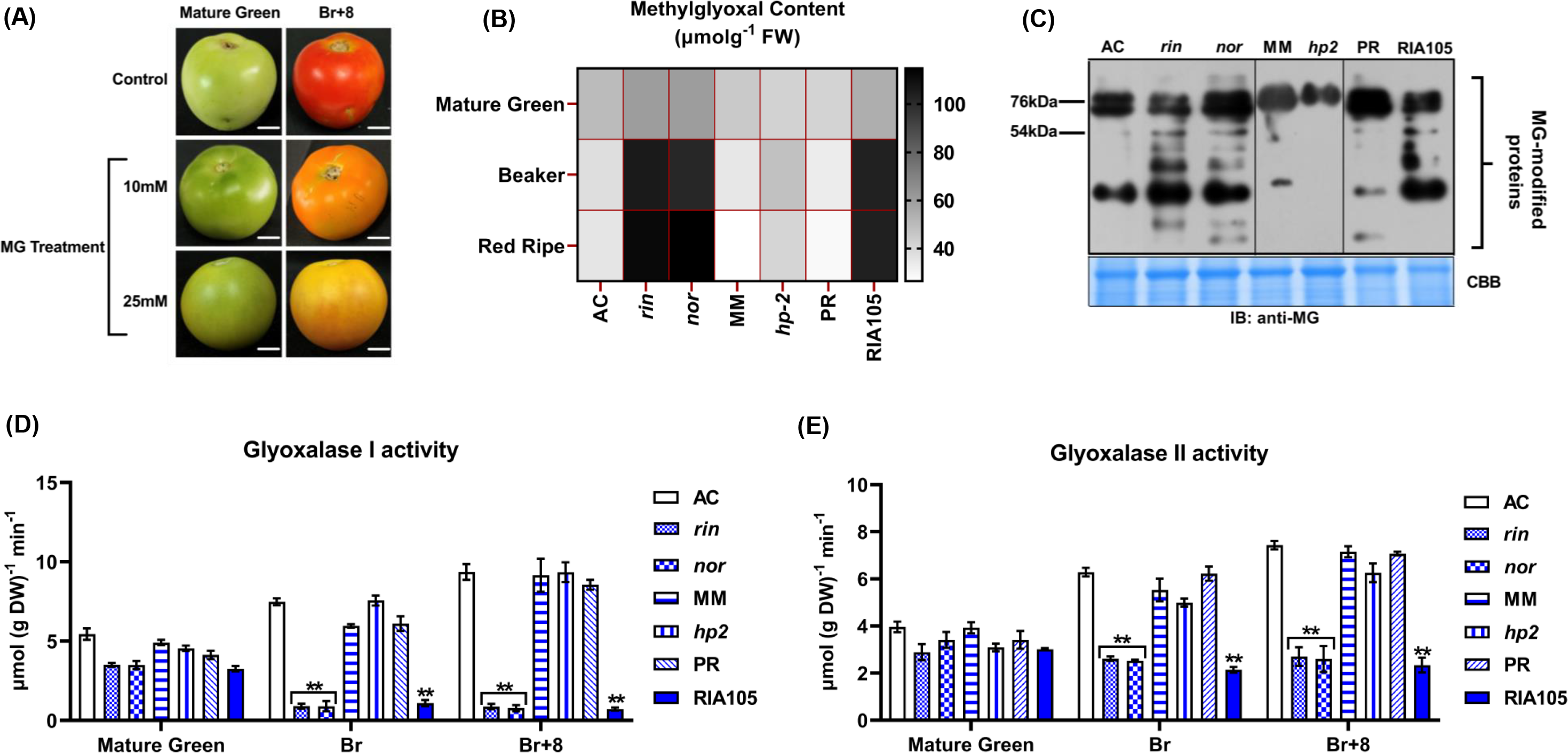
MG serves as a pivotal regulatory factor during tomato fruit ripening. **(A)** Exogenous MG treatment to WT fruits. Mature green fruits of WT (Pusa Ruby background) were injected with a 10 and 25 mM MG aqueous solution. After the treatment, fruits were incubated in a culture room at 26°C, under a 16 h light/8 h dark cycle with a light intensity of 100 μmol m^-2^ s^-1,^ and photographed after seven days.**(B)** Heat-map representation of MG content in WT (Ailsa Craig and Pusa Ruby), *ripening-inhibitor* mutant (*rin*; accession no. LA1795, in the unknown background), and *non-ripening* mutant (*nor*; in Ailsa Craig background) and *35S:RINRNAi* knockdown lines (RIA105 and RIA106; in Pusa Ruby background), *hp2* mutant and its WT (Moneymaker background), at various stages of fruit ripening. PR, Pusa Ruby; MM, Moneymaker. **(C)** Western blot analysis of total proteins to assess the extent of MG-modified protein levels using anti-MG antibodies in different WT (Ailsa Craig, Pusa Ruby, Moneymaker), *rin*, *nor* and *hp2* mutants and *35S:RINRNAi* knockdown lines (RIA105 and RIA106; in Pusa Ruby background) at Br+4 stage. Equal loading is shown using Coomassie brilliant blue (CBB) stained membrane (bottom panel). **(D, E)** Spectrophotometric estimation of enzyme activities of **(D)** Glyoxalase 1 (GLYI) and **(E)** Glyoxalase 2 (GLYII) in different WT, ripening mutant *35S:RINRNAi* knockdown lines at various stages of fruit ripening. Error bars, mean ± SD of three biological replicates. Asterisks indicate the statistical significance using Student’s t-test: *, 0.01 < P-value < 0.05; **, 0.001 < P-value < 0.01, *** p-value < 0.001.

### The glutathione-dependent glyoxalase-detoxification system is responsible for declined MG levels during tomato fruit ripening

The higher levels of MG observed in mature green fruits must result from fruit photosynthesis-related metabolic processes, while at the breaker stage, it could be due to the respiratory burst. We next hypothesized that a glyoxalase detoxification system must be reinforced to lower the MG levels at the onset of ripening to salvage the ripening phenomena from the negative effects of its high concentrations. To substantiate this hypothesis, we performed biochemical assays to measure glyoxalase GLYI and GLYII activity in WTs and corresponding ripening mutants at different stages of ripening (Fig 1D,E). In agreement with the observed MG levels, GLYI and GLYII enzyme activities displayed a ripening-specific pattern, with its catalytic potential increasing as the ripening progressed. In contrast, the enzyme activities were greatly diminished (by almost 80%) in *rin* and *nor* mutant fruits at Br and Br+8 stages relative to control fruits of corresponding genetic backgrounds (Fig 1D,E). Nevertheless, *hp2* genotype fruits represented similar GLYI and GLYII catalytic potential profiles compared to their respective WT fruits (Fig 1D,E). This data signals that activation of the glyoxalase detoxification system is imperative in maintaining a lower MG repertoire, a prerequisite for fruits undergoing a ripening program. The following qRT-PCR analysis of the full complement of tomato glyoxalase I members in WT fruits at the mature green, Br, and Br+8 stages identified *SlGLYI4* and *SlGLYI7* as the members with a characteristic ripening associated expression pattern (Fig 2). *SlGLYI4* showed the strongest induction at the Br and Br+8 stage (10-fold), while *SlGLYI7* transcripts peaked at the Br stage (4-fold) (Fig 2). No significant alteration in the mRNA levels of *SlGLYI5*, *SlGLYI6*, *SlGLYI8*, *SlGLYI9*, and *SlGLYI10* was observed at different ripening stages (Supporting fig S2). On the other hand, *SlGLYI1*, *SlGLYI13,* and *SlGLYI15* showed low transcript levels in all the stages examined (Supporting fig, S2). Within the four-membered SlGLYII subfamily, *SlGLYII3* exhibited no modulations in its expression profile during different stages of ripening, whereas *SlGLYII4* expression declined from mature green to Br+8 stage (Supporting fig, S2). In this analysis, *SlGLYII2* remained the sole member with 3-fold and 5-fold up-regulated transcript levels at the Br and Br+8 stages, respectively (Fig 2). In contrast, the mRNA abundance of SlGLYIII genes (the third subfamily of glyoxalases) remained largely unchanged during fruit ripening, indicating the predominant role of the glutathione-dependent pathway in managing MG levels during fruit ripening (Supporting fig, S2). Taking into account the statistical enrichment of the transcript levels of *SlGLYI4*, *SlGLYI7,* and *SlGLYII2* at Br and Br+8 stages in WT fruit, we then examined the mRNA abundance of these genes in *rin*, *nor* and *hp2* mutant genotypes at the mature green, Br and Br+8 stage (Fig 2). Analogous to the WT fruits, the mRNA accumulation of *SlGLYI7* followed a similar fashion in mutant fruits as that of WT, with peaking at the Br stage (Fig 2). In contrast, approximately 60-70% reduction in the transcript levels of *SlGLYI4* and *SlGLYII2* was observed in *rin* and *nor* fruits at Br and Br+8 stage compared to WT fruits (Fig 2). However, *hp2* mutant fruits displayed no significant modulations in the transcript abundance of *SlGLYI4* and *SlGLYII2* at the onset of ripening when compared to their corresponding WT control (Fig 2). The abovementioned observation signifies that the expression of *SlGLYI4* and *SlGLYII2* in tomato fruits is ripening-regulated. To decipher the functional contribution of *SlGLYI4*, *SlGLYI7,* and *SlGLYII2* in the ripening mechanism, we proceeded with transient silencing of these genes using the VIGS approach. The *TRV2-SlGLYI4* infiltrated fruits revealed mottled green and orange sectors, which diverged from the uniform orange phenotype witnessed in *TRV2* control fruits (Supporting fig, S3). However, *TRV2-SlGLYI7* and *TRV2-SlGLYII2* silenced fruits were devoid of any visual alterations in the pigment accumulation and closely resembled the phenotype of the control *TRV2* fruits (Supporting fig, S3). These results imply that *SlGLYI4* might be an important regulatory component of the MG-detoxification system during tomato fruit ripening.

**Figure 2.**
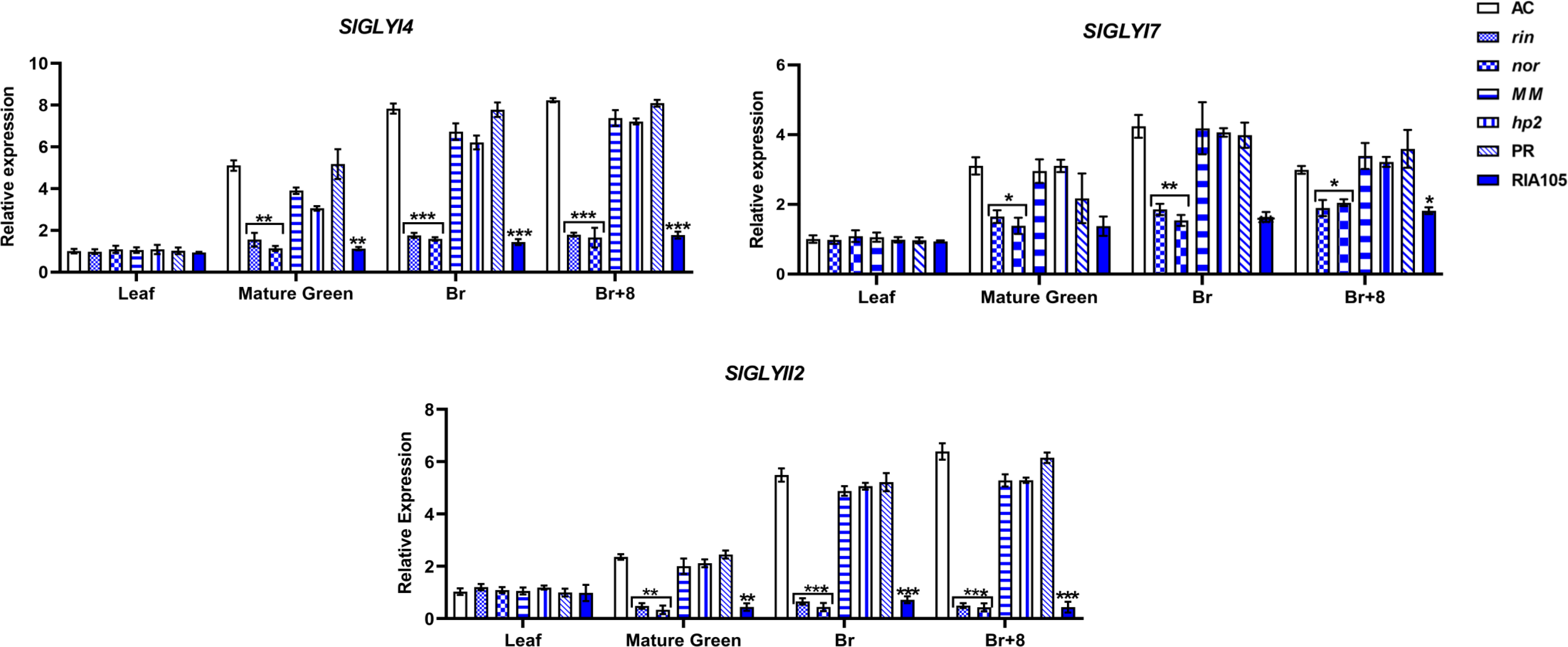
*SlGLYI4* displays a ripening-associated transcript profile in tomato. Expression profiles of *SlGLYI4*, *SlGlYI7,* and *SlGLYII2* in different WT (Ailsa Craig, Pusa Ruby, Moneymaker), *rin*, *nor* and *hp2* mutants and *35S:RINRNAi* knockdown lines (RIA105 and RIA106) at various stages of fruit ripening. PR, Pusa Ruby, AC, Ailsa Craig; MM, Moneymaker. *Actin* was used as the internal control. Error bars, mean ± SD of three biological replicates. Asterisks indicate the statistical significance using Student’s t-test: *, 0.01 < P-value < 0.05; **, 0.001 < P-value < 0.01, *** p-value < 0.001.

### *SlGLYI4* is a direct target of RIN and functions in ethylene dependent manner

The observed activation of *SlGLYI4* during ripening further prompted us to examine its correlation with the gaseous hormone ethylene. For this, we first mined the 2-kb upstream region of *SlGLYI4* for the presence of putative Ethylene Response Elements (ERE) using the PLACE signal search tool (http://www.dna.affrc.go.jp/PLACE/signalscan.html) (Fig 3A). As expected, the promoter sequence of *SlGLYI4* was found to harbour three ERE and two E-box elements, an Ethylene biosynthetic element (LeCPACS2) (Fig 3A). A putative fruit-specific element (TCTTCACA) was also found to be present in this promoter (Fig 3A). We next profiled the transcript abundance of *SlGLYI4* upon treatment with ethylene and its inhibitor, 1-MCP, in Pusa Ruby fruits and noticed a 5-fold induction in its transcripts level in ethylene-treated fruits (Fig 3B). In contrast, a reduction in its mRNA accumulation was observed in 1-MCP treated fruits (Fig 3B). Interestingly, we also noticed the presence of three putative CArG box elements in the retrieved promoter sequence of *SlGLYI4* (Fig 3A). This observation encouraged us to investigate the transcriptional regulation of *SlGLYI4* by RIN, a MADS-box protein known for its ability to bind to CArG boxes in the promoter region of several ripening-associated genes (Fujisawa et al., 2014). The strong activation of *HIS2* reporter gene of the pHIS2.1 *GLYI4_pro_* construct in yeast-one-hybrid (Y1H) assay indicated direct binding of RIN to the promoter of *SlGLYI4* (Fig 3C). Promoters of *SlACS2* was considered positive control for the Y1H experiment, respectively (Fig 3C). To further confirm the interaction of RIN with the promoter of *SlGLYI4 in planta*, we carried out the transient expression assays using GFP reporter system in *N. benthamiana* leaves (Fig 3D). Transactivation of the *SlGLYI4* promoter by RIN significantly enhanced the GFP reporter activity and validated the association between RIN and SlGLYI4 in planta (Fig 3D). Further, the electrophoretic mobility assay (EMSA) provided conclusive evidence of RIN binding to the CArG box in the promoter of *SlGLYI4* (Fig 3E). Together these results implied that RIN promotes the transcriptional activity of *SlGLYI4* by directly binding to its promoter region.

**Figure 3.**
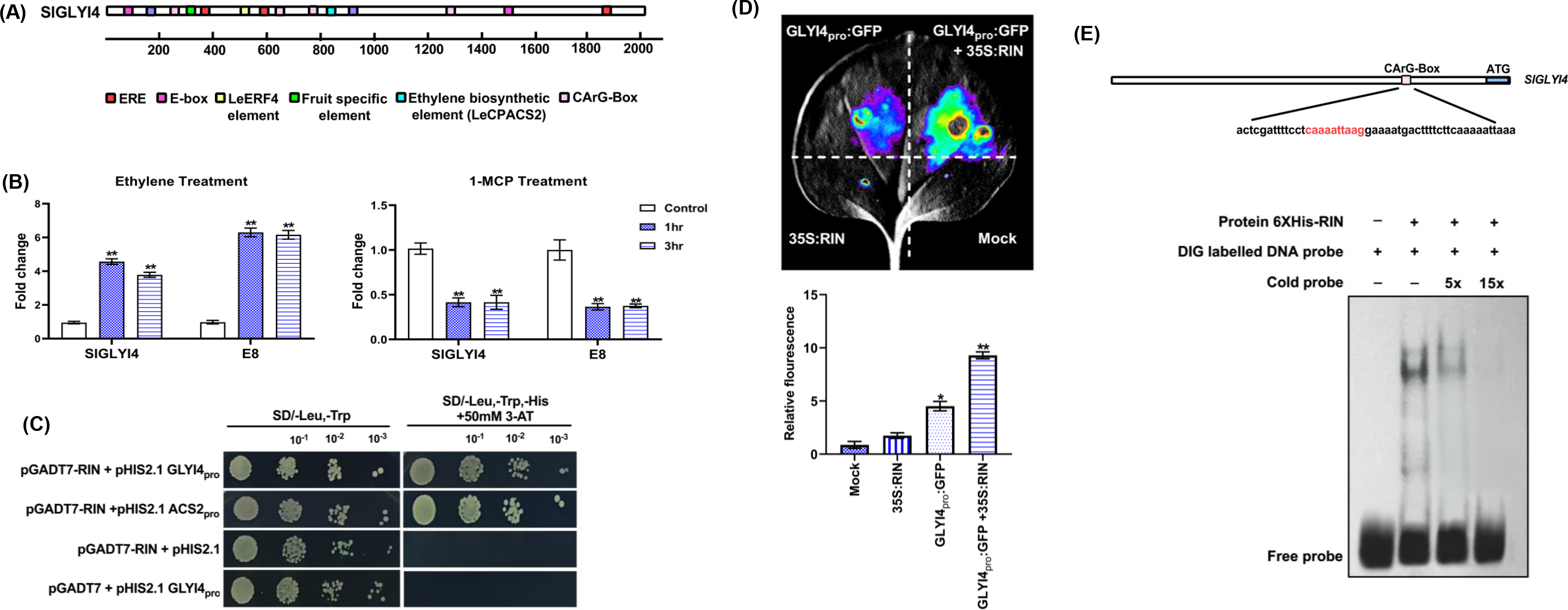
RIN- and ethylene-dependent action of *SlGlYI4* **(A)** 2-kb promoter region of *SlGLYI4* gene analyzed *in silico* for the presence of CArG box, a putative RIN-binding element in addition to other fruit-ripening specific elements. The *cis*-acting regulatory elements identified are represented by different color boxes.**(B)** RT-qPCR analysis of *SlGLYI4* transcripts in total RNA samples extracted from WT mature green fruit samples treated with 100 µM Ethrel and 100 µM 1-MCP. Error bars, mean ± SD of three biological replicates. *E8* was used as a positive control of the treatments. Asterisks indicate the statistical significance using Student’s t-test: *, 0.01 < P-value < 0.05; **, 0.001 < P-value < 0.01, *** p-value < 0.001 . E8, an ethylene response gene.**(C,D)** Identification of DNA binding activity of RIN to the promoter of *SlGLYI4* with **(C)** growth performance of transformants on SD/−Leu−/Trp/−His medium containing 50 mM 3-AT. Binding of RIN to the promoter of tomato *ACC synthase2* (*ACS2*) gene **(D)** *In vivo* interaction study of RIN to promoters of *SlGLYI4* via GFP reporter assays in *N.benthamiana* leaves. Quantitative analysis of fluorescence intensity in three independent determinations was assessed. Error bars represent ±SD of three biological replicates. Asterisks indicate the statistical significance using Student’s t-test: *, 0.01 < P-value < 0.05; **, 0.001 < P-value < 0.01. **(E)** EMSA assay showing RIN binding to the CArG element in the promoter of *SlGLYI4*. The wild type probe containing the CArG motif was digoxigenin-labelled. Competition for SlGLYI4 binding was performed with 5x and 15x cold probes. The symbols - and + represent the absence or presence of the probes and 6X Histidine-tagged RIN protein.

### *SlGLYI4*-silencing significantly alters the fruit’s oxidative landscape

To better understand the molecular mechanism involving *SlGLYI4*-mediated MG homeostasis during ripening, we generated two sets of artificial microRNA-based *SlGLYI4*-silenced transgenic plants; one driven by *CaMV35S* promoter for constitutive silencing and a second set driven by a ripening-specific promoter, *RIP1* (Fig 4A). A total of 10 and 8 independent *SlGLYI4* silencing transgenic lines were obtained for *35S:amiRGLYI4* and *RIP1:amiRGLYI4*, respectively. Two representative lines for each set of constructs i.e., *RIP1:amiRGLYI4#1* and *RIP1:amiRGLYI4#3* for *RIP1:amiRGLYI4*; *35S:amiRGLYI4#5* and *35S:amiRGLYI4#6* for *35S:amiRGLYI4*, were retained for further characterization in T_2_ generation based on the lines showing the best gene silencing of *SlGLYI4*. At the molecular level, *RIP1:amiRGLYI4#1* and *RIP1:amiRGLYI4#3* displayed the strongest suppression of *SlGLYI4* transcript from Br+4 stage onwards, showing only 15-20% accumulation of its levels in WT at the Br+15 stage (Fig 4C). In contrast, *35S:amiRGLYI4#5* and *35S:amiRGLYI4#6* exhibited a severe reduction in *SlGLYI4* expression at all stages, including the mature green stage (Fig 4C). No off-target silencing of other GLYI subfamily members was found in both sets of *SlGLYI4*-silenced lines emphasizing the specificity of the artificial microRNA-based silencing (Supplementary fig, S4). In agreement with the reduction in mRNA levels of *SlGLYI4*, we observed highly inhibited glyoxalase enzymatic activity in *35S:amiRGLYI4* and *RIP1:amiRGLYI4* lines fruits (Fig 4C). The catalytic potential of GLYI enzyme in *RIP1:amiRGLYI4#1* and *RIP1:amiRGLYI4#3* fruits was reduced by 70-75% at the Br+4 stage (Fig 4C). Similarly, *35S:amiRGLYI4#5* and *35S:amiRGLYI4#6* transgenic fruits preserved only 10-15% of GLYI enzymatic activity at all the ripening stages as compared to WT (Fig 4C). Since the loss of GLYI activity is synonymous with elevated MG levels, a 55-60% surge in MG levels was observed in *RIP1:amiRGLYI4#1* and *RIP1:amiRGLYI4#3* transgenic fruit at Br+8 stage in comparison to WT counterparts (Fig 5). A more intense MG accumulation with levels amounting to approximately 70-75% was observed in *35S:amiRGLYI4#5* and *35S:amiRGLYI4#6* fruits at the Br+8 stage relatively to control fruits (Fig 5). Analogously, the repercussions of elevated MG levels in terms of protein modification were more profound in *35S:amiRGLYI4#5* and *35S:amiRGLYI4#6* than their *RIP1:amiRGLYI4* counterparts, as assessed using anti-MG antibodies during different stages of ripening in WT and two sets of transgenics genotypes fruits (Fig 4D). However, *RIP1:amiRGLYI4#1* and *RIP1:amiRGLYI4#3* also exhibited a considerable increase in MG-modified proteins at Br+4 and Br+8 stages compared to WT fruits (Fig 4D). Since MG spontaneously combines with GSH and reduces its availability for other cellular redox reactions, we quantified the reduced GSH content in parallel with measuring the GSH/GSSG ratio in WT and *SlGLY14*-silenced fruits at the onset of ripening (Fig 5). Indeed, a depleted pool of GSH combined with regression in GSH/GSSG ratio was observed in both *35S:amiRGLYI4* and *RIP1:amiRGLYI4* transgenic fruits at late Br stages against the WT fruits (Fig 5). With this shift in the redox state of *SlGLYI4*-silenced transgenic fruits, we next examined the antioxidant reserves in the two sets of *SlGLYI4*-silenced fruits (Fig 5). In harmony with reduced GSH levels, total antioxidant capacity in *35S:amiRGLYI4#5* and *35S:amiRGLYI4#6* was reduced to approximately 20% of WT fruits at the Br+8 stage (Fig 5). On the other hand, *RIP1:amiRGLYI4#1* and *RIP1:amiRGLYI4#3* presented a 65-70% reduction in total antioxidant levels at ripening stages (Fig 5). Altogether, the data implies that the constitutive or ripening-specific silencing of *SlGLYI4* culminates in elevated MG levels in tomato fruit, leading to substantial metabolic and redox modulations inside the fruit, thereby influencing the overall ripening process.

**Figure 4.**
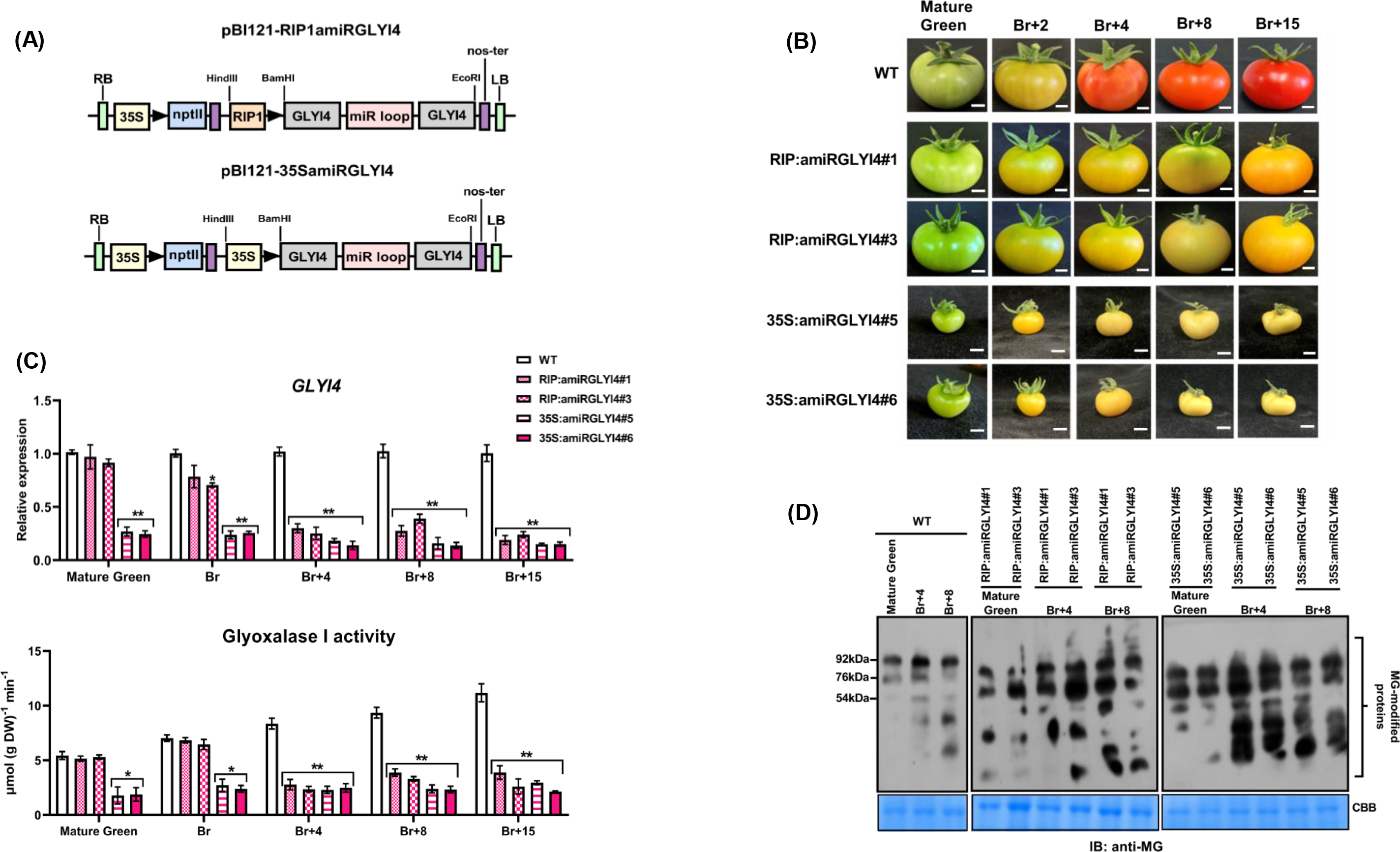
Morphometric analysis and molecular characterization of *35S:amiRGLYI4* and *RIP1:amiRGLYI4* transgenic fruits **(A)** Vector diagram of *pBI121-RIP1::amiRSlGLYI4* (ripening-specific silencing) and *pBI121-35S::amiRSlGLYI4* (constitutive silencing) constructs.**(B)** Phenotypic analysis of WT, *35S:amiRGLYI4* and *RIP1:amiRGLYI4* fruits at mature green (MG), breaker (Br), 4-day after Br (Br+4), 8-day after Br (Br+8), and 15-day after Br (Br+15). **(C)** Transcript levels of *SlGLYI4* in WT, *35S:amiRGLYI4* and *RIP1:amiRGLYI4* transgenic fruit analyzed at mature green (MG), breaker (Br), 3-day after Br (Br+3), 8-day after Br (Br+8), and 15-day after Br (Br+15) by RT-qPCR with *Actin* gene as an internal control. Error bars mean ±SD of three biological replicates. Asterisks indicate the statistical significance using Student’s t-test: *, 0.01 < P-value < 0.05; **, 0.001 < P-value < 0.01. **(D)** Western blot analysis of total proteins to assess the extent of MG-modified protein levels using anti-MG antibodies in WT, *35S:amiRGLYI4,* and *RIP1:amiRGLYI4* transgenic fruits at different stages of fruit ripening. Equal loading is shown using Coomassie brilliant blue (CBB) stained membrane (bottom panel).

**Figure 5.**
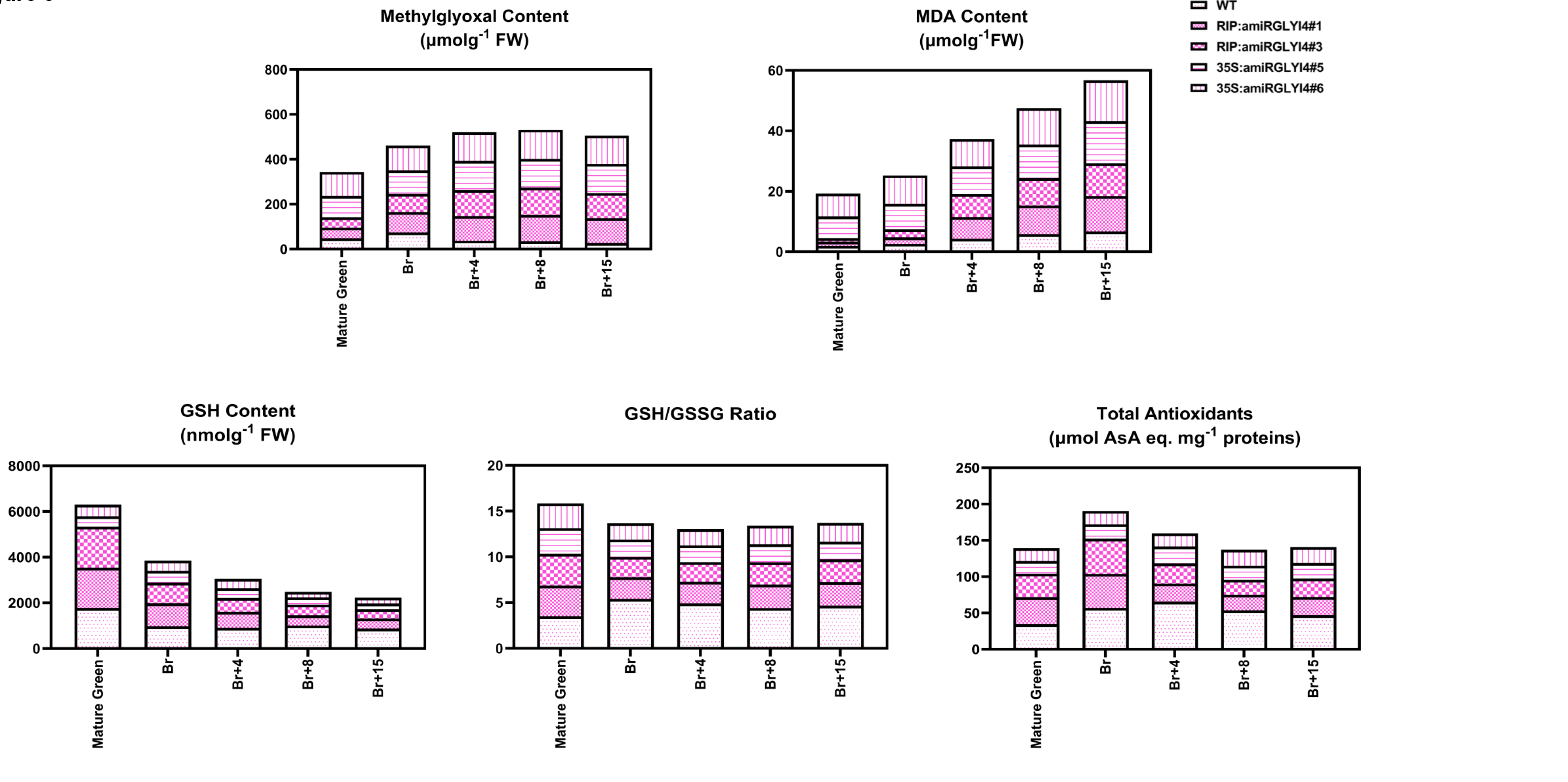
Impact of *S/GLY/4* silencing on the oxidative landscape of transgenic tomato fruits. Spectrophotometric quantification of MG, MDA, reduced glutathione (GSH), GSH/GSSG ratio, and ascorbic acid (AsA) in WT and *35S:amiRGLY/4* and *RIP1:amiRGLY/4* transgenic fruits at mature green (MG), breaker (Br), 4-day after Br (Br+4), 8-day after Br (Br+8), and 15-day after Br (Br+15) stages of fruit ripening.

### Loss of *SlGLYI4* directly impacts fruit pigmentation and texture during ripening

Apart from the observed off-balanced redox state of the transgenic fruits, other cardinal outcomes of an aggravated MG accumulation embodied alterations in fruit color development and texture. Phenotypically, fruits from both sets of transgenic lines strongly mimicked the yellow-colored phenotype of MG-infiltrated fruits at the Br+8 stage, which persisted till the later stages of ripening (Fig 4B). In conjugation with alterations in pigment accumulation, *35S:amiRGLYI4#5* and *35S:amiRGLYI4#6* transgenic plants displayed a drastic reduction in fruit size compared to WT and *RIP1:amiRGLYI4* genotypes, possibly accounting for the fact that glyoxalases have been designated as markers for cell growth and differentiation (Fig 4B). HPLC-based lycopene and β-carotene profiling in WT and transgenic pericarp tissues at the mature green, Br, Br+4, Br+8, and Br+15 stages revealed a 65-70% reduction in lycopene accumulation at the Br+4 stage of *RIP1:amiRGLYI4#1* and *RIP1:amiRGLYI4#3* fruit tissue (Fig 6A). The *35S:amiRGLYI4#5* and *35S:amiRGLYI4#6* transgenic fruits showed even more drastic inhibition in pigmentation and accumulated 70-80% less lycopene than their WT counterparts (Fig 6A). Concomitantly, we noticed a sharp rise in β-carotene levels in *RIP1:amiRGLYI4* and *35S:amiRGLYI4* transgenic pericarp tissues at the Br+15 stage, perfectly resonating with the yellow color of these fruits (Fig 6A). On par with a more severe reduction in lycopene amount, *35S:amiRGLYI4#5* and *35S:amiRGLYI4#6* line fruits displayed a 3-fold excess of β-carotene concentration at Br+8 stage relative to WT fruits (Fig 6A). To uncover the molecular basis responsible for these skewed pigment profiles of both sets of *SlGLYI4*-silenced lines, we then analyzed the expression pattern of genes involved in carotenoid biosynthetic pathways at different stages of ripening (Fig 6B). The qRT-PCR analysis showed a profound decrease in mRNA levels of a key regulatory gene of the carotenoid biosynthesis pathway, *PSY1* (*Phytoene synthase1*), in both sets of transgenic fruits than WT at Br+4 and Br+8 stages (Fig 6B). An equivalent reduction of approximately 70-80% in transcript abundance of *PDS* (*Phytoene desaturase*) was observed in both sets of *SlGLYI4*-silenced genotype fruits than WT fruits at the Br+8 stage (Fig 6B). The expression profiles of *ZDS (Carotene desaturase)* and *CRTISO (Carotenoid isomerase)* displayed similar impairment in their transcript levels in the two sets of transgenic fruits at the Br+8 stage compared to control fruits (Fig 6B). However, *35S:amiRGLYI4* line fruits presented more intense repression of these transcripts at all the examined stages than their *RIP1:amiRGLYI4* counterparts (Fig 6B). In contrast, transcripts of *LCY-B (Lycopene cyclase*) and *LCY-E*, the genes mediating β-carotene accumulation in fruits, showed a substantial upregulation in their mRNA abundance at Br+4 and Br+8 stages in *RIP1:amiRGLYI4* and *35S:amiRGLYI4* fruits against the control fruits (Fig 6B). This data indicates that enhanced MG concentration in tomato fruits results in a modified lycopene to β-carotene ratio due to alterations in the transcript patterns of key carotenoid pathway genes.

**Figure 6.**
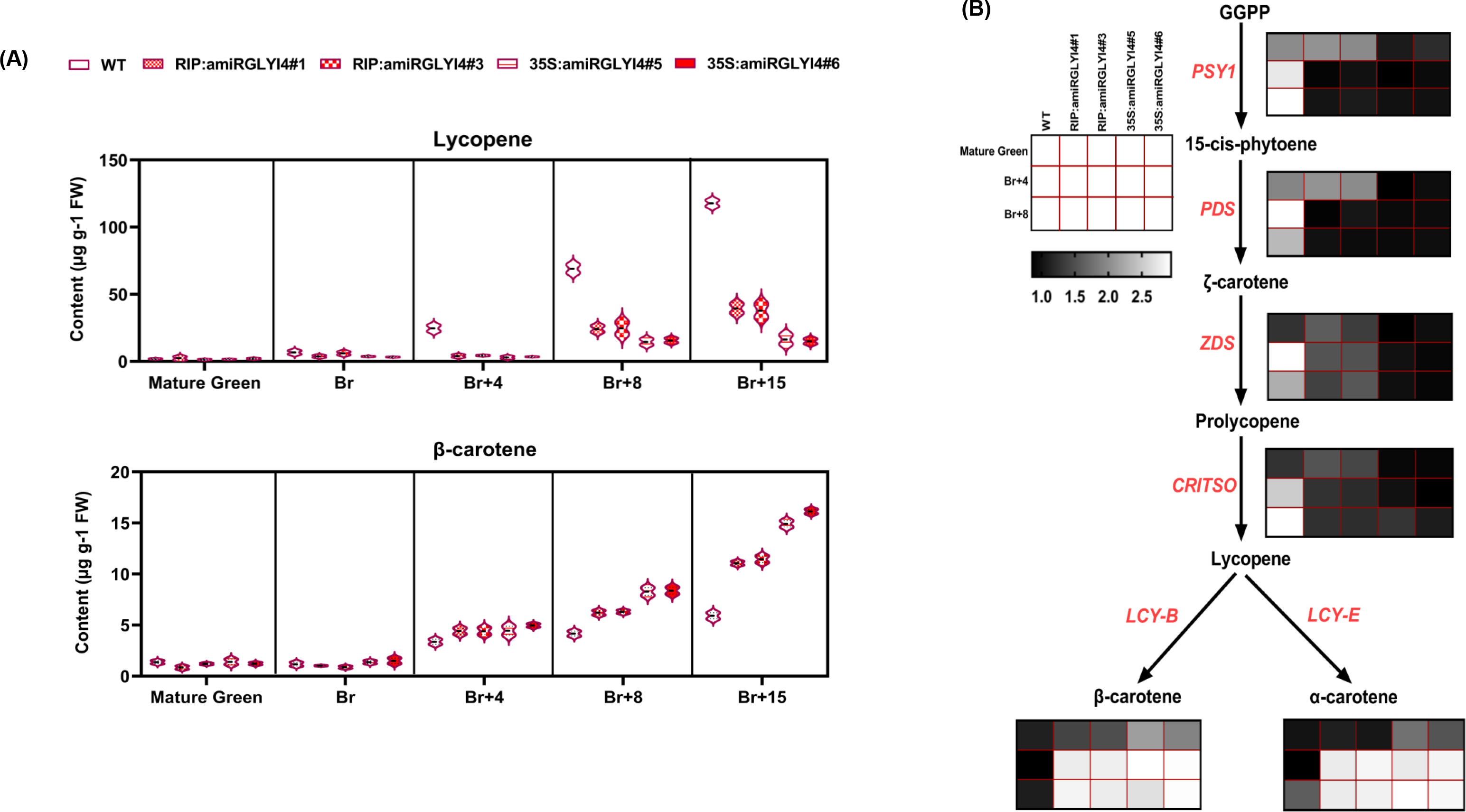
Assessment of pigment accumulation in S/GLY/4-silenced fruits **(A)** Box plot representation of HPLC-based estimation of lycopene and -carotene in WT and *35S:amiRGLY/4* and *RIP1:amiRGLY/4* transgenic fruits at different stages of fruit ripening. **(B)** Heat map representation of RT-qPCR analysis of carotenoid biosynthetic pathway genes in WT and *35S:amiRGLY/4* and *RIP1:amiRGLY/4* transgenic fruits at mature green (MG), 4-day after Br (Br+4) and 8-day after Br (Br+8) with *Actin* gene as an internal control. The scale for each heat map is provided. PSY1 phytoene synthase; PDS phytoene desaturase; ZDS, carotenoid desaturase; CRTISO carotenoid isomerase; LCYE: lycopene epsilon cyclase; LCYB: lycopene beta cyclase

Since the outer pericarp is essential in determining the rate of expansion and mechanical support for the whole fruit, we next investigated the texture of the pericarp of both sets of *SlGLYI4*-silenced lines. Fruits from *RIP1:amiRGLYI4#1* and *RIP1:amiRGLYI4#3* transgenic lines had significantly firmer outer pericarp tissues at Br+8 and Br+15 stages compared to the WT fruits (Fig 7A). As observed earlier for other traits, *35S:amiRGLYI4#5* and *35S:amiRGLYI4#6* fruits overpowered their *RIP1:amiRGLYI4* equivalents displaying almost 70% de-acceleration in terms of the degree of softening (Fig 7A). We then examined the mRNA abundance of various critical cell wall modifying enzymes such as *PL (Pectate lyase)*, *PG (Polygalactouronase)*, *PE (Pectin esterase)*, *TBG4 (Tomato beta-galactosidase 4)* using qRT-PCR in WT and transgenic genotypes at ripening stages. *35S:amiRGLYI4#5* and *35S:amiRGLYI4#6* fruits accumulated low levels of *PL* and *PG* transcripts at the Br+8 stage compared to control fruits (Fig 7B). The characteristic ripening-induced expression of *PL* and *PG* could not be observed at the onset of maturation in *RIP1:amiRGLYI4#1* and *RIP1:amiRGLYI4#3* fruits (Fig 7B). Likewise, a 60-65% reduction was witnessed in *TBG4* and *PE* transcripts in *RIP1:amiRGLYI4* and *35S:amiRGLYI4* transgenic plants at the Br+8 stage (Fig 7B). The centrality of pectin modifications during fruit softening has long been recognized. Keeping that in mind, we next estimated the PL activity, formulated on its capacity to catalyze a β-eliminative reaction with the cell-bound pectin in WT and transgenic lines at the Br+8 stage. Synchronous to the expression profiles of *PL*, the enzymatic potential of PL was significantly reduced in *SlGLYI4* silenced fruits relative to the control ones (Fig 7D). To substantiate these findings, we measured the pectin content of pectin at the Br+8 stage in the transgenic and control fruits (Fig 7C). Consistent with the curtailed PL activity in tandem with increased fruit firmness, *RIP1:amiRGLYI4#1* and *RIP1:amiRGLYI4#3* transgenic fruits accumulated enhanced amounts of proto-pectin whereas the soluble form of pectin was decreased by about 30% in comparison to the WT fruits (Fig 7C). Likewise, proto-pectin levels elevated by 25%, whereas soluble pectin fell by approximately 35-40% in *35S:amiRGLYI4#5* and *35S:amiRGLYI4#6* fruits relative to the control ones at the Br+8 stage (Fig 7C). Altogether, these results signify the adverse effects of *SlGLYI4*-silencing and enhanced levels of MG on cell wall depolymerization, a quintessential process necessary for the fruit to become palatable.

**Figure 7.**
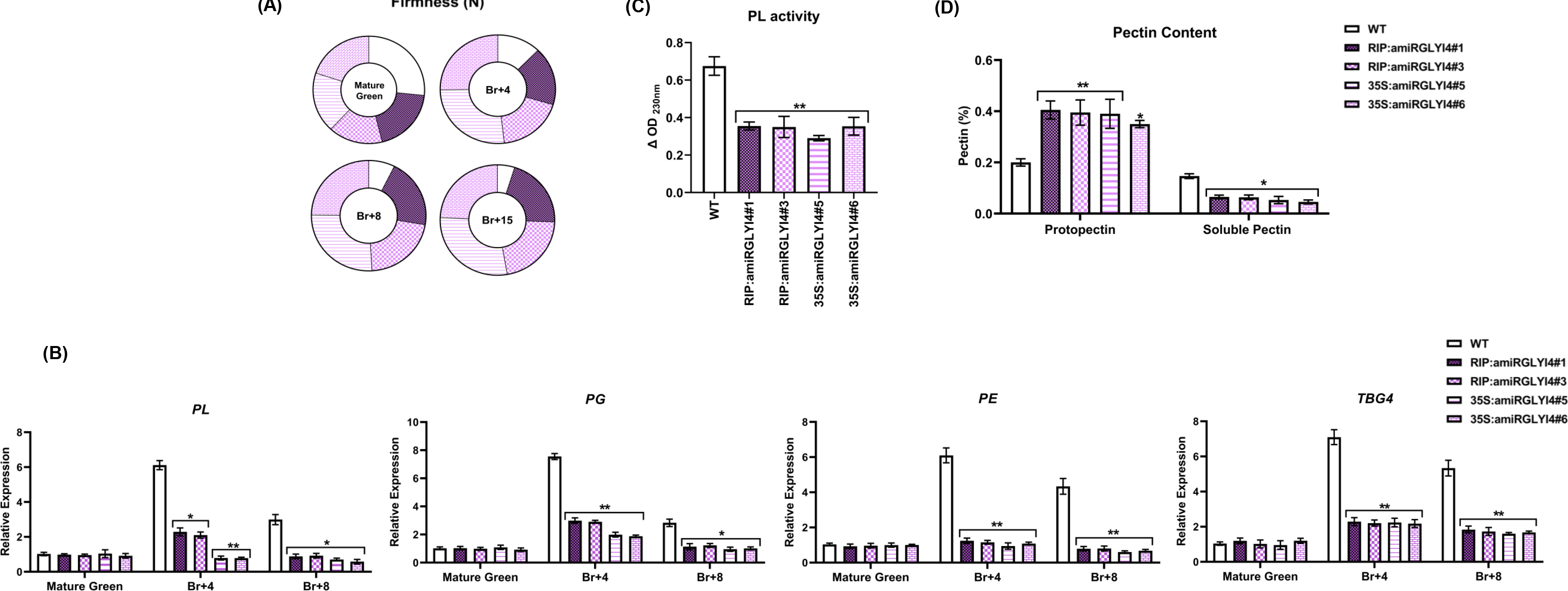
Modulations in levels of fruit firmness in *35S:amiRGLYI4* and *RIP1:amiRGLYI4* transgenic fruits **(A)** Fruit firmness analysis in WT, *SlGLYI4*-silenced line fruits at different stages of ripening. A total of 10 fruits were used for each measurement, and the values shown are the means ±SD. **(B)** RT-qPCR analysis of pectate lyase (*PL*), polygalacturonase (*PG*), pectinesterase (*PE*), and β-galactosidase (*TBG4*) at mature green (MG), 4-day after Br (Br+4), 8-day after Br (Br+8) in *35S:amiRGLYI4* and *RIP1:amiRGLYI4* and WT fruits. *Actin* was used as the internal control. The Error bars represent ±SD of three biological replicates. Asterisks indicate the statistical significance using Student’s t-test: *, 0.01 < P-value < 0.05; **, 0.001 < P-value < 0.01. **(C)** Assessment of pectate lyase enzyme activity in WT, *35S:amiRGLYI4* and *RIP1:amiRGLYI4* silenced fruits at 8-day after Br (Br+8). The Error bars represent ±SD of three biological replicates. Asterisks indicate the statistical significance using Student’s t-test: *, 0.01 < P-value < 0.05; **, 0.001 < P-value < 0.01. **(D)** Estimation of pectin level in terms of protopectin and soluble pectin in WT and *SlGLYI4*-silenced fruits at Br+8. The Error bars represent ±SD of three biological replicates. Asterisks indicate the statistical significance using Student’s t-test: *, 0.01 < P-value < 0.05; **, 0.001 < P-value < 0.01.

### The key ripening regulators are differentially expressed in *SlGLYI4*- silenced fruits

The observed ramifications on the color and texture of *35S:amiRGLYI4* and *RIP1:amiRGLYI4* fruits prompted us to perform the transcript profiling of major fruit ripening regulators. Compared to the control fruits, the mRNA levels of *RIN (Ripening-inhibitor)*, *CNR (Colorless non-ripening),* and *NOR* (*Non-ripening*) were astonishingly low at Br+8 and Br+15 stages in *35S:amiRGLYI4* and *RIP1:amiRGLYI4* fruits (Fig 8). Moreover, the transcriptional regulators of pigment synthesis, i.e., *FUL1* and *FUL2*, in both sets of *SlGLYI4*-silenced fruits, revealed a 60-70% reduction in transcript accumulation at the Br+8 stage against the WT fruits (Fig 8). Further, *TAGL1*, another MADS-box ripening regulator, showed a similar reduction in its mRNA abundance in *35S:amiRGLYI4* and *RIP1:amiRGLYI4* fruits at Br+8 and Br+15 stages when compared to WT fruits (Fig 8).

**Figure 8.**
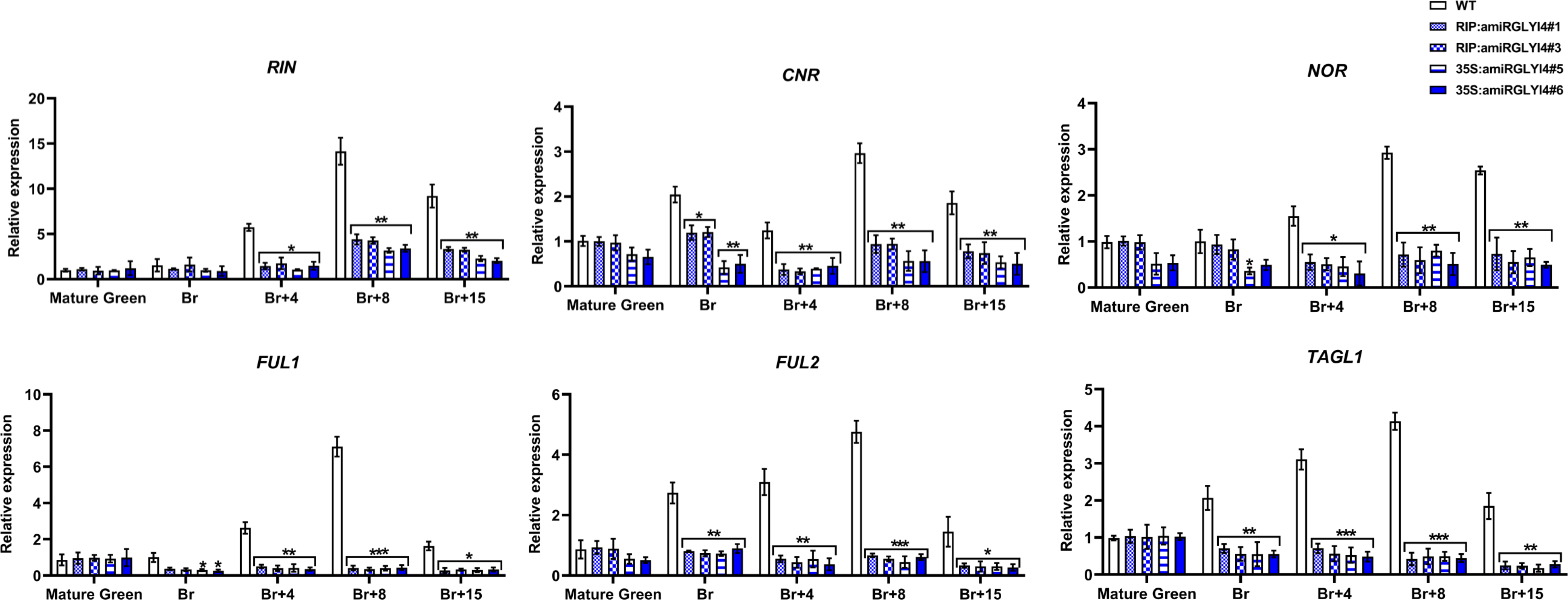
Transcript profiling of key ripening regulator genes in WT, *35S:amiRGLYI4* and *RIP1:amiRGLYI4* silenced tomato lines during fruit ripening. Total RNA was extracted from mature green (MG), breaker (Br), 4-day after Br (Br+4), 8-day after Br (Br+8), and 15-day after Br (Br+15) stages of *SlGLYI4-* silenced and WT fruits. The relative mRNA levels of each gene were normalized using the *Actin* gene as an internal control. Error bars represent ±SD of three biological replicates. Asterisks indicate the statistical significance using Student’s t-test: *, 0.01 < P-value < 0.05; **, 0.001 < P-value < 0.01. RIN ripening inhibitor; NOR non-ripening; CNR colorless non-ripening; TAGL1 tomato AGAMOUS-LIKE 1; FUL1 and FUL2 Fruitful MADS-box transcription factor homologs.

### MG overaccumulation inhibits ethylene biosynthesis in *SlGLYI4*-silenced fruits

Enhanced ethylene production at the onset of the Br stage is instrumental in galvanizing ripening-associated modulation in pigment synthesis and cell wall metabolism, a hallmark of all climacteric fruits. To check if ethylene biosynthesis is perturbed due to MG overaccumulation, we quantified its levels at different stages of ripening in transgenic and non-transgenic fruits (Fig 9A). Unlike the sharp classical increase in ethylene levels from mature green to Br stage in WT, *35S:amiRGLYI4#5* and *35S:amiRGLYI4#6* fruits failed to attain such robust autocatalytic activation of ethylene at Br and Br+4 stages (Fig 9A). Among the two sets of transgenic plants, *RIP1:amiRGLYI4#1* and *RIP1:amiRGLYI4#3* fruits showed a transitory increase in ethylene production from mature green to Br stage, which was followed by a sharp decline from Br to Br+4 stage (Fig 9A). As expected, no such transitory increase in ethylene biosynthesis was observed in *35S:amiRGLYI4* fruits (Fig 9A). To connect these alterations to the genetic level, assessment of *ACS2*, *ACS4* (*1- aminocyclopropane-1-carboxylate synthase 4*), and *ACO1* (*1- aminocyclopropane-1-carboxylate oxidase 1*) transcripts were carried out in control and the two sets of transgenic fruits during ripening (Fig 9B). In compliance with the shift in ethylene profile, both *RIP1:amiRGLYI4* and *35S:amiRGLYI4* fruits showed reduced mRNA levels of the ethylene biosynthetic genes at late Br stages (Fig 9B). No significant alterations were noted in the transcript profiles of the ethylene receptor genes i.e., *ETR2*, *ETR3*, *ETR4,* and *ETR5* and ethylene signaling positive regulators such as *EIN2* and *EIL2* in WT and *SlGLYI4*-silenced fruits (Fig 9B). However, on analyzing the endogenous transcript levels of ripening-related ERFs, a 60-70% downregulation in the expression levels of key ethylene response factors, with a known role in fruit ripening, such as *SlERF.E1*, *SlERF.E2* and *SlERF.E4* were detected in *RIP1:amiRGLYI4* and *35S:amiRGLYI4* fruits at late Br stages in contrast to the WT counterparts (Fig 9B). Furthermore, to decipher a link between the elevated MG accumulation and inhibited ethylene production, exogenous ethylene was supplied to both sets of *SlGLYI4*-silenced line fruits at the mature green stage. Astonishingly, the fruits of *RIP1:amiRGLYI4#3* and *35S:amiRGLYI4#5* transgenic plants treated with ethylene reversed the non-ripening phenotype with visual enhancement in lycopene production at 4-days post-Br stage (Fig 9C). Overall, these results suggest that the observed ripening defects in *SlGLYI4*-silenced fruits are due to inhibited ethylene production.

**Figure 9.**
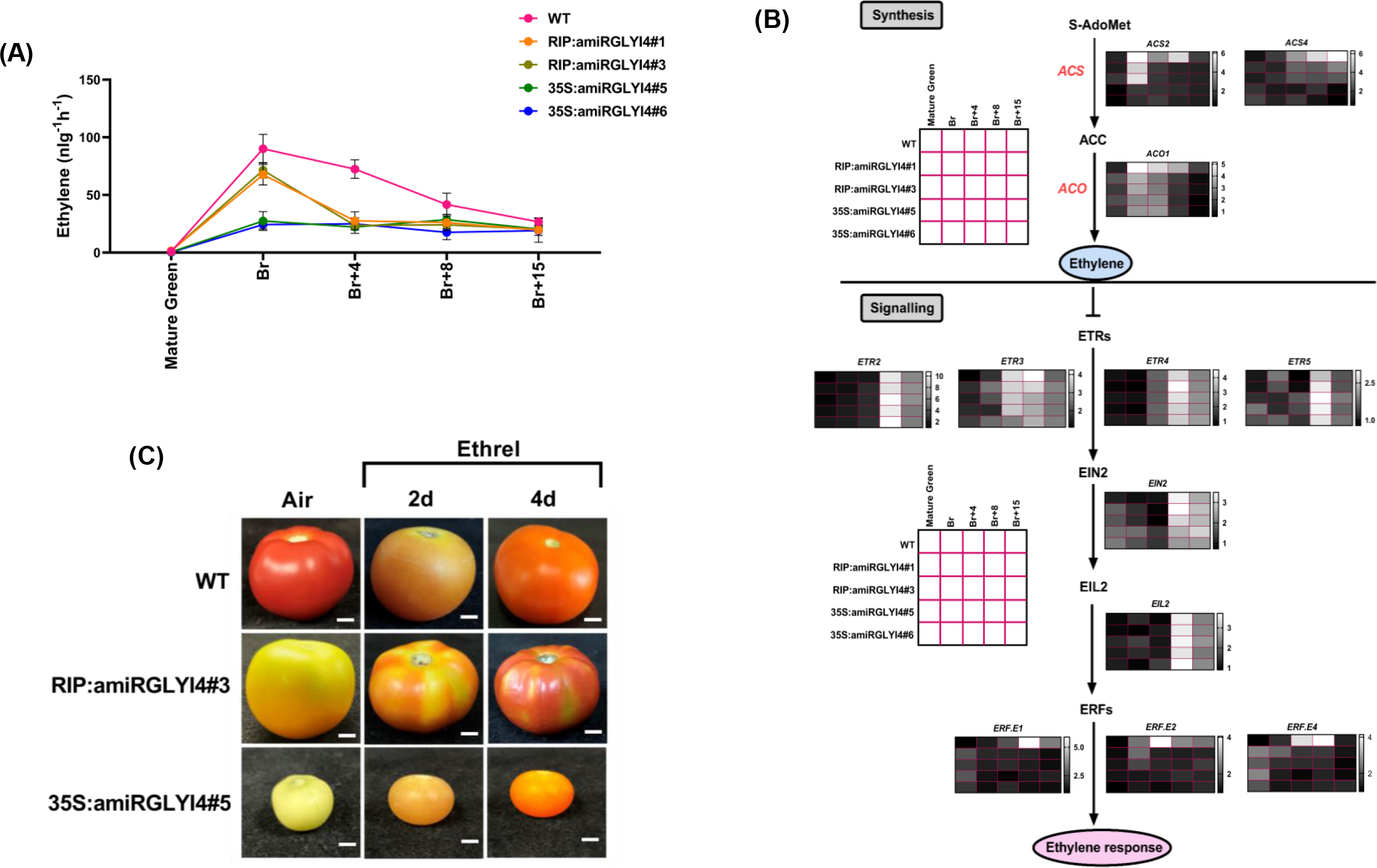
Alterations in ethylene biosynthesis and perception in *SlGLYI4-*silenced transgenic fruits. **(A**) Ethylene production of WT, *35S:amiRGLYI4* and *RIP1:amiRGLYI4* silenced fruits assessed at mature green (MG), breaker (Br), 4-day after Br (Br+4), 7-day after Br (Br+8), and 15-day after Br (Br+15) stages. Values represent the means of at least five individual fruits. The Error bars represent ±SD of three biological replicates. **(B)** Heat map representation of RT-qPCR analysis of ethylene biosynthesis and perception pathway genes at mature green (MG), breaker (Br), 4-day after Br (Br+4), 8-day after Br (Br+8), and 15-day after Br (Br+15) in WT, *35S:amiRGLYI4* and *RIP1:amiRGLYI4* silenced fruits with *Actin* gene as an internal control. The scale for each heat map is provided individually. ACO1, aminocyclopropane-1-carboxylic acid oxidase 1; ACS2 and ACS4 aminocyclopropane-1-carboxylic acid synthases; ETR2, ETR3, ETR4, and ETR5 ethylene receptors; EIN2 ethylene signalling protein; EIL2 EIN2-like protein; Ethylene response factors ERF.E1, ERF.E2 and ERF.E4. **(C)** Exogenous ethylene treatment on WT and *SlGLYI4* silenced fruits. Mature green fruits from WT, *35S:amiRGLYI4,* and *RIP1:amiRGLYI4* transgenic lines were injected with a buffer solution containing 10 mM MES, pH 5.6, sorbitol (3% w/v) and 100 µM Ethrel (2-Chloroethylphosphonic Acid, 40% Solution, SRL Diagnostics). Control fruits were injected only with the buffer. After the treatment, fruits were incubated in a culture room at 26°C, under a 16 h light/8 h dark cycle with a light intensity of 100 μmol m^-2^ s^-1,^ and photographed after four days.

### Free soluble methionine content is reduced in *SlGLYI4*-silenced fruits

To uncover the possible cause of the incompetence of *RIP1:amiRGLYI4* and *35S:amiRGLYI4* fruits to synthesize climacteric endogenous ethylene, UPLC-based detection of methionine, the precursor of ethylene, was carried out in WT and *SlGLYI4-*silenced fruits at different stages of ripening. To our surprise, *35S:amiRGLYI4#5* and *35S:amiRGLYI4#6* fruits exhibited the maximum reduction of approximately 50% and 60-70% in methionine amount at Br and Br+8 stages, respectively, compared to WT fruits (Fig 10). Likewise, *RIP1:amiRGLYI4#1* and *RIP1:amiRGLYI4#3* transgenic fruits accumulated significantly reduced methionine at late maturation stages compared to WT controls (Fig 10). Since methionine is designated as a member of the aspartate pathway family, we next examined the levels of other amino acids of this family, aspartate, lysine, and isoleucine, and found that their levels remained largely unaltered at the Br+4 and Br+8 stages of ripening in all three genotypes (Fig 10). However, *RIP1:amiRGLYI4* and *35S:amiRGLYI4* fruits presented a statistically significant reduction of about 20-25% in threonine content at the Br+4 stage than the WT control fruits (Fig 10). To further substantiate the non-ripening phenotype of *SlGLYI4*-silenced fruits due to low methionine input verging on impaired ethylene output, *RIP1:amiRGLYI4#3* and *35S:amiRGLYI4#5* mature green fruits were externally supplemented with 10 mM methionine (Fig 11A). Surprisingly, this treatment failed to restore the normal fruit pigmentation in *SlGLYI4*-silenced fruits, as evident by no signs of lycopene accumulation in these genotypes at the Br+4 stage as compared to methionine-treated WT fruits (Fig 11A). Additionally, the compensatory application of methionine to these fruits was ineffective in catalyzing ethylene production or lycopene accumulation, evident by no substantial increase in either of the parameters at the Br+4 stage (Fig 11C,D). This observatio was further supported at the genetic level by the unaltered transcript levels of *ACS2* and *ACS4* genes in methionine-treated *RIP1:amiRGLYI4#3* and *35S:amiRGLYI4#5* versus the only buffer-treated control fruits (Fig 11D). Altogether, these results indicate that although *SlGLYI4*-silenced fruits accumulated inhibited methionine profile during the ripening process, the ethylene biosynthetic pathway downstream of methionine may also be adversely impacted due to high MG levels in *SlGLYI4*-silenced genotypes.

**Figure 10.**
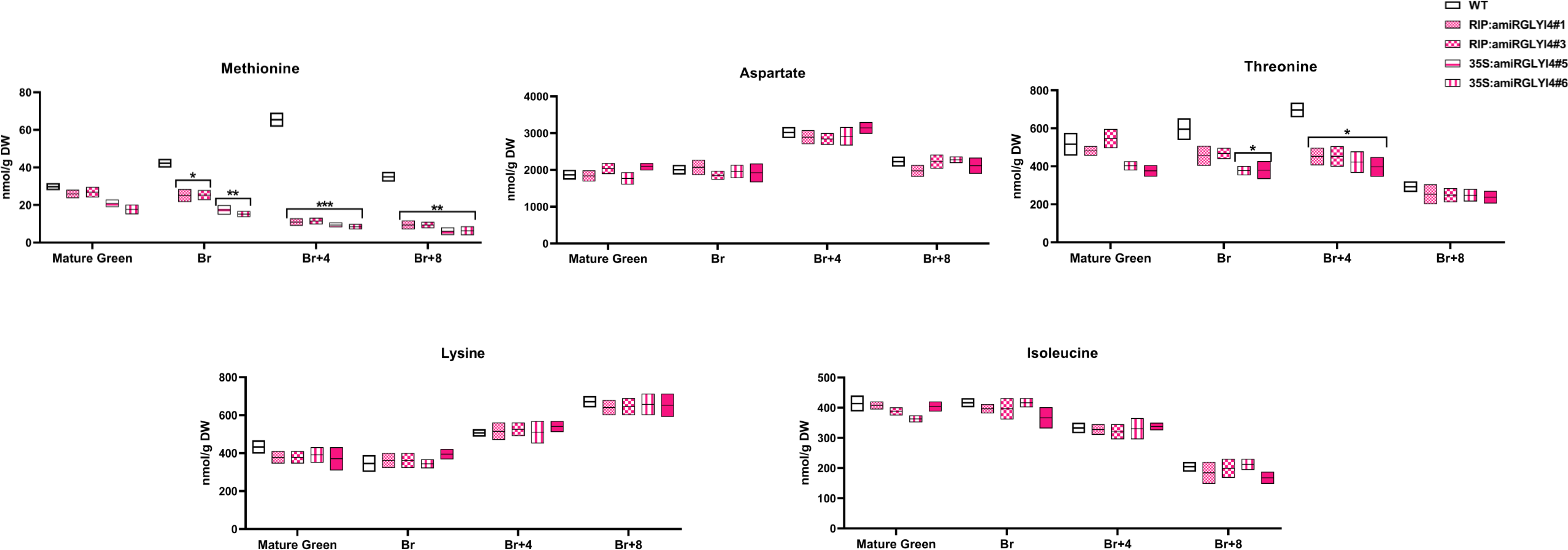
Free soluble amino acid profiling in S/GLY/4-silenced transgenic fruits. Box plot representing the content of Aspartate pathway family of amino acids in WT, *35S:amiRGLY/4* and *RIP1:amiRGLY/4* fruits at mature green (MG), breaker (Br), 4-day after Br (Br+4) and 8-day after Br (Br+8). Values represent the means of at least five individual fruits. The Error bars represent ±SD of three biological replicates. Asterisks indicate the statistical significance using Student’s t-test: *, 0.01 < P-value < 0.05; **, 0.001 < P-value < 0.01.

**Figure 11.**
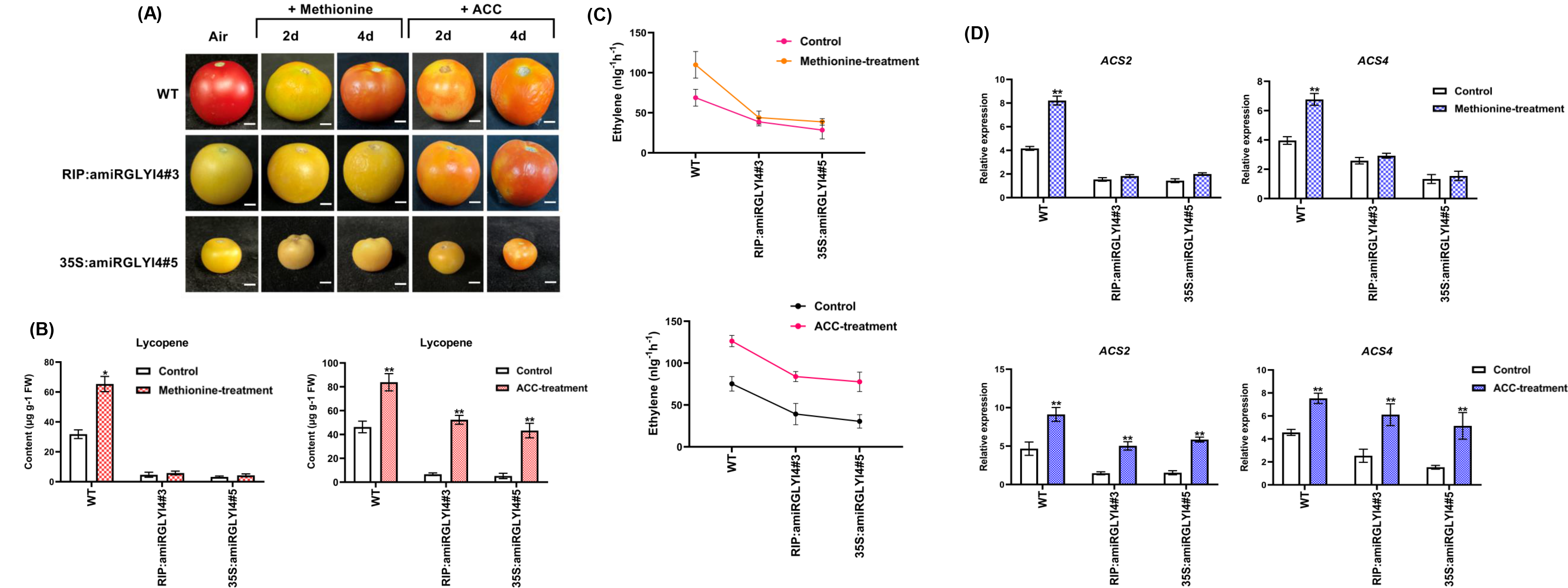
Methionine and ACC feeding to *SlGLYI4*-silenced transgenic fruits **(A**) Exogenous methionine and ACC treatment on WT and *35S:amiRGLYI4* and *RIP1:amiRGLYI4* fruits. Mature green fruits from WT and *SlGLYI-4* silenced lines were injected with 10mM methionine and 100µM ACC. After the treatment, fruits were incubated in a culture room at 26°C, under a 16 h light/8 h dark cycle with a light intensity of 100 μmol m^-2^ s^-1,^ and photographed after four days. **(B)** HPLC-based lycopene estimation post methionine and ACC treatment in WT (WT), *35S:amiRGLYI4* and *RIP1:amiRGLYI4* fruits at 4-day after Br (Br+4). Values represent the means of at least five individual fruits. The Error bars represent ±SD of three biological replicates. Asterisks indicate the statistical significance using Student’s t-test: *, 0.01 < P-value < 0.05; **, 0.001 < P-value < 0.01. **(C)** Ethylene production of post methionine and ACC treatment in WT, *35S:amiRGLYI4* and *RIP1:amiRGLYI4* fruits at 4-day after Br (Br+4). Values represent the means of at least five individual fruits. The Error bars represent ±SD of three biological replicates. **(D)** Transcript profiling of ethylene biosynthetic genes, *ACS2* and *ACS4*, in WT, *35S:amiRGLYI4* and *RIP1:amiRGLYI4* transgenic fruits at 4-day after Br (Br+4). Values represent the means of at least five individual fruits. The relative mRNA levels of each gene were normalized using the *Actin* gene as an internal control. The Error bars represent ±SD of three biological replicates. Asterisks indicate the statistical significance using Student’s t-test: *, 0.01 < P-value < 0.05; **, 0.001 < P-value < 0.01.

### ACC feeding of *SlGLYI4*-silenced fruits reversed their non-ripening phenotype

The contrasting phenotypes of ethylene- and methionine-treated *RIP1:amiRGLYI4#3* and *35S:amiRGLYI4#5* fruits prompted us to investigate further the negative effects of the enhanced MG accumulation on other components of the ethylene biosynthetic pathway. Apart from methionine, the application of a more direct precursor of ethylene, ACC (1-aminocyclopropane-1-carboxylic acid), to *RIP1:amiRGLYI4#3* and *35S:amiRGLYI4#5* mature green fruits activated the ethylene production as apparent by a 40-45% rise in ethylene levels in these fruits at Br+4 stage in comparison to the buffer-treated control fruits (Fig 11C). These results were strongly supported at the molecular level by the elevated transcript levels of *ACS2* (50%) and *ACS4* (30%) in ACC-treated *RIP1:amiRGLYI4#3* fruits than their buffer-treated controls at the Br+4 stage (Fig 11D). A similar increase in the mRNA abundance of both the genes at Br+4 was observed in *35S:amiRGLYI4#5* genotype fruits (Fig 11D). Phenotypically, ACC-treated *RIP1:amiRGLYI4#3* and *35S:amiRGLYI4#5* fruits attained the characteristic orange to the red color associated with the tomato fruits undergoing the normal ripening process (Fig 11A). To corroborate these results quantitatively, we next conducted HPLC-based lycopene measurement assays in ACC-treated and control *RIP1:amiRGLYI4#3* and *35S:amiRGLYI4#5* fruits at Br+4 stage (Fig 11B). In comparison to the control fruits, *RIP1:amiRGLYI4#*3 and *35S:amiRGLYI4#5* fruits exhibited a 50-60% enrichment in lycopene levels in these fruits (Fig 11B). This data concludes that besides displaying alterations in methionine content, a build-up of MG in *SlGLYI4*-silenced fruits directly impacts ethylene biosynthesis at multiple levels, including ACC.

## Discussion

Patterns of MG accumulation in plants are strongly influenced by photosynthesis-dependent sugar metabolism. Therefore, its effective elimination via the glyoxalase detoxification pathway is essential in maintaining MG level to prevent cellular toxicity (Shimakawa *et al*., 2014; Sankaranarayanan *et al*., 2015). The basic catalytic module of the glyoxalase detoxification system is bienzymatic, in which GLYI enzyme-mediated reaction is designated as the rate-limiting step of the pathway (Sankaranarayanan *et al*., 2015; Li, 2016; Rabbani, 2022). Much of our understanding of how valuable the glyoxalase system is to the growth and development of plants has come from the studies on model organisms, Arabidopsis, and rice, often based on the propensity of cellular damage associated with elevated MG accumulation. In Arabidopsis, a reduction in triose phosphate isomerase activity involved in the glycolytic pathway, resulted in abnormally high levels of MG that triggered inhibition of the post-germinative establishment of seedlings (Chen and Thelen, 2010). On similar terms knock out of a rice glyoxalase gene, *OsGLYI3*, has been implicated in decreasing seed longevity and increasing sensitivity to salt stress (Liu *et al*., 2022). Apart from being labelled as MG detoxifier, a glyoxalase 1 protein has been annotated as a compatibility factor, the degradation of which leads to self-incompatibility response in *Brassica napus* (Sankaranarayanan *et al*., 2015). Manipulation of the glyoxalase enzymatic potential in several plant species has been reported to impart stress tolerance against multiple abiotic and biotic stresses (Madan *et al*., 2005; Yadav *et al*., 2005; Bhomkar *et al*., 2008; Álvarez Viveros *et al*., 2013). Though GLYI enzymes have been reported to regulate various physiological and developmental processes, amazingly, their role remained largely uninvestigated in fruit ripening.

This study reports a downward trend in MG accumulation in ripening fruits. The pericarp tissue of mature green tomato fruits, a photosynthetically active stage, accumulates maximum levels of MG that gradually declines with the progression of ripening. This resonates perfectly well with reports documenting increased MG production in photosynthetically active green tissues of plants (Phillips and Thornalley, 1993; Hoque *et al*., 2016*b*; Singla-Pareek *et al*., 2020). As the chloroplast degrades with the onset of ripening, the increased metabolic activities associated with the respiratory burst might lead to enhanced levels of MG. However, the MG levels were curtailed during later stages of ripening which synchronizes well with the observed escalation in the GLYI and GLYII detoxification potentials in the wild-type genotype as the ripening progressed. In addition, the exogenous application of MG could de-accelerate tomato fruit ripening, as indicated by significantly low levels of lycopene accumulation in these fruits. The impact of altered MG homeostasis on the fruit maturation process was reinforced with the defective ripening mutants, *rin,* and *nor* exhibiting enormous levels of MG at late Br stages. The disturbed MG equilibrium was found to be in harmony with decreased GLYI and GLYII activities that were sufficient to alter the extent of protein modifications by MG in *rin* and *nor* mutant fruits. Thus, the perturbed MG equilibrium appears to be a conserved response in several non-ripening mutants of tomato. These results point toward a central role of MG homeostasis as one of the critical factors regulating tomato fruit ripening.

Because of a strong up-regulation in the endogenous transcript levels at different ripening stages with a strong suppression in *rin* and *nor* mutant fruits, *SlGLYI4* could be regarded as a ripening-specific regulator responsible for maintaining low MG levels at the initiation of ripening. A relatable enhancement in GLY1 protein levels to mitigate the soaring amounts of MG in the papillary cells for successful pollination was documented in *Brassica napus* (Sankaranarayanan *et al*., 2015). The responsiveness of *SlGLYI4* to external ethylene application in tomato mature green fruits further synchronizes with its ripening-associated profile. In this study, we demonstrated that *SlGLYI4* is under the direct regulation of LeMADS-RIN (RIN), as substantiated by the binding and activation of *SlGLYI4* promoter by RIN using Y1H (heterologous system), GFP transactivation (in planta) and EMSA (in-vitro) assays. RIN is characterized as one of the global developmental regulators of ripening by virtue of its ability to directly target key-ripening related genes (Vrebalov *et al*., 2002; Li *et al*., 2019). The capacity of RIN to increase the transcriptional activity of *SlGLYI4* by binding to its promoter is in agreement with the down- regulation of *SlGLYI4* transcripts abundance in *rin* mutant fruits. A different data set supporting the role of *SlGLYI4* in controlling the level of MG and, in turn, fruit ripening was generated through detailed analysis of artificial microRNA-based silencing lines of *SlGLYI4* under constitutive (*CaMV35S*) and ripening specific (*RIP*) promoters (Aggarwal *et al*., 2017). The diminished MG detoxification potential of both sets of *SlGLYI4*-silenced genotypes was in alignment with the significant enhancement of MG concentration in the transgenic fruits at different stages of ripening. Since the extent of MG-modified proteins was remarkably increased in both sets of *SlGLYI4*-silenced fruits, the defect in the normal ripening process could be attributed to the glycation of single or multiple proteins crucial for the ripening process as one of the possible consequences of elevated MG levels.

A major outcome of *SlGLYI4* knockdown observed in *35S:amiRGLYI4* fruits was a drastic reduction in the fruit size, whereas the fruits of *RIP1:amiRGLYI4* transgenic plants were comparable in size to the WT fruits. This could be attributed to the constitutive silencing of *SlGLYI4*, resulting in decreased accumulation of GLY1 enzyme and inhibited cell division during the early stages of fruit development. Many reports in both plant and animal systems have confirmed a positive correlation between glyoxalase activity and cell division (Sankaranarayanan *et al*., 2017; Morgenstern *et al*., 2020; Aragonès *et al*., 2021). In Datura callus, an increase in the enzymatic potential of GLYI proteins was concerted with enhanced cell growth, protein, and DNA synthesis, whereas a severe impairment in its activity was observed upon the addition of mitotic inhibitors in the growth media (Ramaswamy *et al*., 1984). Likewise, another study on proliferative cell suspension cultures of *Glycine max L.* reported elevated glyoxalase activity during the logarithmic growth phase with a subsequent decline upon inhibition of cell division, endorsing glyoxalases as markers for cell division (Paulus *et al*., 1993; Sankaranarayanan *et al*., 2017).

Another profound ramification of the enhanced MG accumulation due to *SlGLYI4* silencing in both sets of transgenic genotypes was witnessed on fruit pigment during ripening. Phenotypically, *35S:amiRGLYI4* and *RIP1:amiRGLYI4* yielded yellow-colored fruits with no visual signs of lycopene production even at 15 days post breaker. This incompetency of *SlGLYI4*- silenced fruits to attain the characteristic red color could be attributed to the modified lycopene to the β-carotene ratio in these fruits, as evidenced by the HPLC-based estimation of these pigments at different stages of ripening. The observed off-balance in pigment synthesis might be attributed to a reduction in the expression profile of *PSY1* and *PDS* with a concomitant increase in the transcript levels of *β-LYC* and ε*-LYC* cyclases observed in this study. Several pieces of evidence support the supremacy of *PSY1* and *PDS* genes in signalling lycopene production, whereas the *LYC-B* and *LYC-E* cyclases are the primary targets for enhanced β-carotene concentration in tomato fruits (Giuliano *et al*., 1993; Fray and Grierson, 1993; Fraser *et al*., 1994; Pecker *et al*., 1996; Ronen *et al*., 2000; Fantini *et al*., 2013; Liu *et al*., 2014). The alarmingly low amounts of ethylene production in conjunction with an acute down-regulation of important ripening regulators such as RIN, NOR, FUL1, and FUL2 in *35S:amiRGLYI4* and *RIP1:amiRGLYI4* fruits might have also contributed to reduced lycopene accumulation in these fruits. This was in harmony with the results of the previous experiments demonstrating that RNAi-mediated RIN silencing resulted in yellow-pigmented fruits with complete inhibition of ripening (Vrebalov *et al*., 2002, 2009; Alba *et al*., 2005). Along similar lines, the dual suppression of FUL1 and FUL2 in tomato produced fruits with decreased lycopene accumulation (Bemer *et al*., 2012).

Besides the modulation in pigment synthesis, the extent of fruit softening was also impacted in *SlGLYI4*-silenced fruits. A drastic reduction in the endogenous transcript profiles of genes encoding cell wall modifying enzymes such as *PL*, *PG*, *PE,* and *TBG4* possibly contributed to the enhanced degree of fruit firmness. In the classical ripening literature, *PL*, *PE,* and *PG* are considered major players involved in the cell wall degradation process (Hall *et al*., 1993; Grierson *et al*., 1993; Yang *et al*., 2017). Together with alterations in the severe impairment in the catalytic potential of PL and a surge in amounts of protopectin in *SlGLYI4*-silenced fruits were as per the observed fruit firmness of *35S:amiRGLYI4* and *RIP1:amiRGLYI4* genotypes. Recently, CRISPR generated knockout of *PL* in tomato resulted in firmer fruits with increased shelf life (Wang *et al*., 2019). Also, it has been previously documented that fruit softening is sensitive to ethylene levels (Alba *et al*., 2005; Li *et al*., 2019). In this regard, a defect in ethylene biosynthesis in *SlGLYI4*-silenced fruits could have resulted in failed catalysis of the disintegration of cell wall components.

One of the most significant adverse effects of *SlGLYI4* silencing was observed in the extent of ethylene production in transgenic fruits. The failure of *35S:amiRGLYI4* and *RIP1:amiRGLYI4* fruits to achieve autocatalytic activation of ethylene was in compliance with the decreased expression profiles of *ACS2*, *ACS4,* and *ACO1* genes. A complete cessation of ripening has previously been reported in fruits displaying similar defects in the ethylene biosynthetic pathway at the level of *ACS* and *ACO* genes (Hamilton *et al*., 1990, 1991; Oeller *et al*., 1991). On supplying ethylene externally, the reversal of the non-ripening phenotype of *SlGLYI4*-silenced fruits reiterated probable faults in ethylene production rather than its signaling, a hypothesis validated by unaltered mRNA levels of ethylene receptor genes. Since exogenous ethylene treatment could compensate for low ethylene and carotenoid levels, we needed to measure the amounts of its biological precursor, methionine, in the *SlGLYI4*-silenced transgenic fruits. Recently, the Methionine synthase 2 (MS2) protein, an enzyme catalyzing the synthesis of methionine in Arabidopsis, has been reported as a target of glycation by carbonyl compounds such as glyoxals and MG in response to drought stress (Chaplin *et al*., 2019). Based on this finding, a decline in free soluble methionine content in *35S:amiRGLYI4* and *RIP1:amiRGLYI4* fruits could be attributed to potential protein modification of tomato MS homologs in these genotypes. If the ethylene biosynthesis was impacted only at its precursor level, the methionine feeding of *SlGLYI4*-silenced fruits should have restored the normal ripening. However, external methionine supplementation proved unsuccessful in elevating ethylene and lycopene levels, indicating that high accumulation of MG has impacted ethylene biosynthesis at multiple levels. This hypothesis was further supported by the reversal of non-ripening phenotype of *35S:amiRGLYI4* and *RIP1:amiRGLYI4* fruits upon treatment with ACC, demonstrating an orange fruit coloration at Br+4 stage of ripening. Hence, besides the detected defect in methionine content in *SlGLYI4*- silenced fruits, another enzyme of the Yang’s cycle, catalyzing a step between methionine and ACC synthesis, might be a target of protein modification by increased MG accumulation in these fruits. This was in line with identification of another protein, Methionine Adenosyltransferase 4, susceptible to glycation by MG in response to various stress conditions in Arabidopsis (Chaplin *et al*., 2019). Methionine adenosyl transferase, also known as SAM synthetases (SAMS), is known to catalyze the conversion of methionine to S-adenosylmethionine, the precursor of ACC. SAM and ACC are regarded as the most dedicated regulators of ethylene biosynthesis (Barry *et al*., 2000; Pattyn *et al*., 2021). Extensive analysis has revealed that ethylene production is inextricably linked to the homeostasis of its general precursor S-adenosyl-l-methionine (SAM), which is controlled by transcriptional and posttranslational mechanisms, as well as the metabolic flux through the adjacent Yang cycle (Peleman *et al*., 1989; Pattyn *et al*., 2021). On the other hand, ACC is a unique precursor of ethylene biosynthesis, and recent findings indicate that this small cyclopropane is a signalling molecule independent of ethylene (Van de Poel and Van Der Straeten, 2014; Polko and Kieber, 2019). In this regard, it may be concluded that any possible alteration in the homeostasis of SAM or ACC because of enhanced MG accumulation in *SlGLYI4*-silenced fruits might have contributed to the incompetency of these fruits to undergo the normal ripening process.

In summary, in the present study, we conclusively demonstrate through pharmacological, biochemical, morpho-physiological and genetic tools that declining MG levels during fruit maturation are critical for tomato fruits to undertake a successful ripening program. Out of the GSH-independent and GSH-dependent MG-detoxification pathways, the first system appears to play a predominant role in facilitating fruit transition at the onset of ripening, primarily through *SlGLYI4*-mediated reduction of MG levels. Without *GLYI4* activation during ripening, fruits MG levels drastically increase and target the ethylene biosynthesis pathway, plausibly glycating MS2 and SAM synthase homologs (Fig 12). Although low ethylene levels along with reduced expression of key ripening regulators in *GLYI4*-silenced fruits justify the perturbed pigmentation and fruit shelf-life, a disruption of carotenoid biosynthesis and pectin metabolism through direct glycation of some of the key enzymes of these pathways by the enhanced intracellular MG levels can not be ruled out. Overall, the results presented here underpin a *SlGLYI4-*mediated novel regulatory mechanism of MG detoxification crucial for regulating fruit maturation and ripening program in tomato.

**Figure 12.**
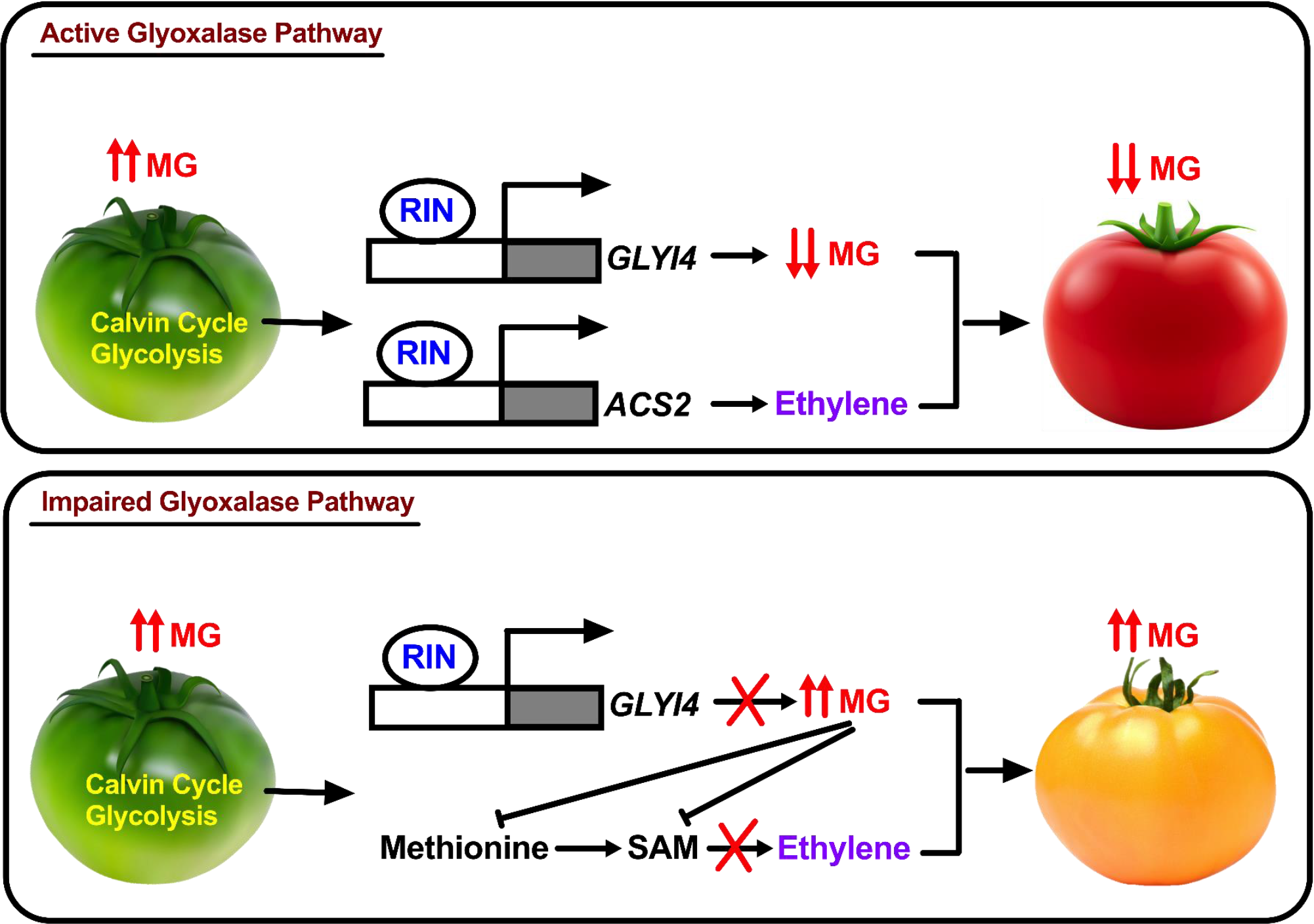
Proposed model depicting the function of glyoxalase detoxification pathway and the impact of MG levels during tomato fruit ripening. The photosynthetically active mature green fruits accumulate high amounts of MG. However, at the onset of ripening, RIN-mediated activation of *SlGLYI4* curtails the enhanced levels of MG associated with the respiratory burst and several other metabolic activities so that the ripening can proceed in a normal way. In the absence of the glyoxalase detoxification system at the time of fruit maturation, soaring amounts of MG, plausibly generated as a consequence of respiratory burst, can potentially inhibit the ethylene biosynthetic pathway at multiple levels, thereby leading to fruits incompetency to ripen. ACS2 1-aminocyclopropane-1-carboxylate synthase 2; SAM S-adenosyl methionine.

## Supporting Data

Table S1 List of primers used in this study.

Table S2 Segregation analysis of kanamycin resistance in T2 progeny of both sets of SlGLYI4-silenced transgenic lines.

Figure S1 On-vine and off-wine treatment of MG on wild-type tomato fruits.

Figure S2 Expression profiling of glyoxalase genes during tomato fruit ripening.

Figure S3 Virus-induced gene silencing of *SlGLYI4*, *SlGLYI7,* and *SlGLYII2* in WT fruits.

Figure S4 Transcript profiling of *GLYI* sub-family members in *SlGLYI4*- silenced lines during different stages of ripening.

## Author contributions

Conceptualization- A.K.S., R.K., S.K.S; Investigation, validation, methodology, and formal analysis- P.G., U.R., A.P., S.K., S.S.; Writing original draft-P.G.’ R.K.; Reviewing and editing- A.K.S., R.K., S.K.S.; Supervision- A.K.S., R.K.

## Accession Numbers

Sequence data from this article can be found in the GenBank/EMBL data libraries under accession numbers. *SlGLYI4* (Solyc02g080630)*, SlGLYI7* (Solyc05g041200)*, SlGLYII2* (Solyc06g053310)*,SlERF*.*E1* (Solyc09g075420), *SlERF*.*E2*(Solyc06g063070), *SlERF*.*E4* (Solyc01g065980), *PSY1* (Solyc03g031860), *PDS* (Solyc03g123760), *ZDS* (Solyc01g097810), *LYC-B* (Solyc04g040190), *LCY-E* (Solyc12g008980), *CRTISO* (Solyc10g081650), *ACS2* (Solyc01g095080), *ACS4* (Solyc05g050010), *ACO1* (Solyc07g049530), *E4* (Solyc03g111720), *E8* (Solyc09g089580), *PG* (Solyc10g080210), *RIN* (Soly c05g012020), *CNR* (Solyc02g077920), *NOR* (Solyc10g006880), *TAGL1* (Solyc 07g055920), *EIN2* (Solyc09g007870), *EIL2* (Solyc01g009170), *EIL3* (Solyc01g 096810), *ETR2* (Solyc07g056580), *ETR3* (Solyc09g075440), *ETR4* (Solyc06g0 53710), *ETR5* (Solyc11g006180),*FUL1* (Solyc06g069430), *FUL2* (Solyc03g11 4830), *PL* (Solyc03g111690)), *PE* (Solyc07g064180), *CRITSO* (Solyc10g081650), *TBG4* (Solyc12g008840).

## Abbreviations

MG: Methylglyoxal
1-MCP: 1- Methylcyclopropane
VIGS: Virus-induced gene silencing
ROS: Reactive oxygen species
RNS: Reactive nitrogen species
GSH: Glutathione
GLY: Glyoxalase

